# The PENGUIN approach to reconstruct protein interactions at enhancer-promoter regions and its application to prostate cancer

**DOI:** 10.1101/2022.10.20.512998

**Authors:** Alexandros Armaos, François Serra, Iker Núñez-Carpintero, Ji-Heui Seo, Sylvan C. Baca, Stefano Gustincich, Alfonso Valencia, Matthew L. Freedman, Davide Cirillo, Claudia Giambartolomei, Gian Gaetano Tartaglia

## Abstract

Here we introduce Promoter-ENhancer-GUided Interaction Networks (PENGUIN), a method to uncover protein-protein interaction (PPI) networks at enhancer-promoter contacts. By integrating H3K27ac-HiChIP data and tissue-specific PPI information, PENGUIN enables cluster enhancers-promoter PPI networks (EPINs) and pinpoint actionable factors.

Validating PENGUIN in cancer (LNCaP) and benign (LHSAR) prostate cell lines, we observed distinct CTCF-enriched clusters, which identifies diverse chromatin conformations. In LNCaP, we found an EPIN cluster enriched with oncogenes and prostate cancer-associated SNPs. We uncovered a total of 208 SNPs in LNCaP EPINs and used CRISPR/Cas9 knockout and RNAi screens to confirm their relevance.

PENGUIN’s application in prostate cancer demonstrates its potential for studying human diseases. The approach allows exploration in different cell types and combinations of GWAS data, offering promising avenues for future investigations. In conclusion, PENGUIN provides valuable insights into the interplay between enhancer-promoter interactions and PPI networks, facilitating the identification of relevant genes and potential intervention targets.

## Introduction

Enhancer-promoter (E-P) interactions play a crucial role in orchestrating gene expression and ensuring the proper regulation of cellular processes. DNA-binding proteins (DBPs), including transcription factors (TFs), act as key players in this regulatory network by binding to enhancers and bridging additional protein interactions between enhancers and promoters. In this work we define Enhancers-Promoter protein-protein Interaction Network (EPIN) as the local interactome connecting a single promoter with all its interacting enhancers. EPIN interactions are facilitated by various types of intermediate proteins, such as co-activators (e.g., mediators), chromatin structural proteins (e.g., cohesin), and noncoding RNA-binding proteins.

While protein-protein interactions (PPIs) have been extensively studied ^1, 2^, the integration of chromatin architecture information, specifically through chromosome conformation capture (3C-like) techniques, with PPI analysis is still in its early stages. Joint investigations of chromatin loops and PPIs are crucial for prioritizing functional interactions ^3^. However, it is important to note that many of these studies often lack the necessary biological context at various levels.

As of today, the characterization of context specific intermediate PPIs involved in disease pathways and their association with DBPs remains largely unanswered ^4^. Previous studies have highlighted the significance of disrupted E-P loops in several human disorders ^5–7^. In cancer, enhancers are frequently subject to sequence and structural variations, leading to the dysregulation of TFs and chromatin modifiers, which contribute to oncogenesis ^8^. Consequently, targeting these enhancer-driven mechanisms holds great promise for therapeutic interventions in cases such as Prostate Cancer (PrCa) ^9^. In this context, advanced techniques such as HiC and its derivative HiChIP ^10^, in combination with ChIP-seq, could enable the identification and characterization of specific chromatin interactions between enhancers and promoters. In particular, H3K27ac-HiChIP has emerged as a powerful tool designed to detect and amplify E-P interactions and has been successfully employed to uncover susceptibility genes associated with cancer, including PrCa ^11^.

To characterize protein interactions that take place at the E-P contacts, we developed the Promoter-ENhancer-GUided Interaction Networks (PENGUIN) approach. For each promoter annotated in the genome and covered by at least one HiChIP interaction, PENGUIN builds an EPIN by integrating several sources of information: (1) high-resolution chromatin interaction maps enriched for a marker of active E-P activity (H3K27ac-HiChiP); (2) tissue-specific physical nuclear PPIs; (3) high-quality curated binding motifs of protein-DNA interactions; (4) tissue specific gene expression, used as a filter of protein data.

To prove the usefulness of our PENGUIN approach, we applied it to uncover EPINs in a PrCA cell line, androgen-sensitive human prostate adenocarcinoma cells (LNCaP), and validate our findings in comparison to a benign prostate epithelial cell line (LHSAR). PrCa is the 2nd most common cancer in men ^12^. Its distinct hormone-dependent nature is characterized by high expression and frequent genetic amplification of *AR*. AR is a regulator of homeostasis and proteases transcription, such as *KLK3* encoding PSA (Prostate-Specific Antigen). *AR* gene is also a principal therapeutically targeted oncogene in PrCa ^13^. Increased genetic instability resulting in chromosomal rearrangements and high frequency of mutations are deemed indicative of PrCa aggressiveness ^14^ for which there is need of *ad hoc* treatments ^15^. Recurrent mutations in *FOXA1*, involved in prostate organogenesis and regulator of *AR* transcription, have been observed in several populations ^16, 17^. Hundreds of PrCa-associated single nucleotide polymorphisms (SNPs) have been identified by genome-wide association studies (GWAS), including genomic regions within tumor suppressor genes and oncogenes, such as *MYC* ^18^. However, the functional relationship between most of these SNPs and PrCa pathophysiology is unknown. This missing part of the picture, together with the growing evidence of abnormal transcriptional programs driven by genetic instability, led us to investigate the role of chromatin architecture in PrCa. In particular, we focused on the nuclear proteins potentially involved in transcriptional regulation through the interaction of promoters and non-coding regulatory elements, enhancers.

By clustering together promoters with similar EPIN structures, PENGUIN identified 273 promoters whose genes are enriched in PrCa fine-mapped SNPs, known PrCa oncogenes, and ChIP-Seq-validated binding sites of transcriptional repressor CTCF. The proteins that populate such EPINs constitute putative PrCa-related factors, some of which have not been previously described to be associated with PrCa SNPs or oncogenes. Moreover, the EPINs detected by PENGUIN enable the characterization of distinct molecular cascades enriched in PrCa SNPs at E-P contacts. These represent new potential molecular targets in PrCa that cannot be identified through conventional analytical procedures, such as E-P contacts and GWAS overlap. To explore our results we made a dedicated server available at https://penguin.life.bsc.es/.

Our methodology, focussing at the specific EPIN resolution level, reveals a new relation between 3D genome conformation and disease phenotype. This new relation allows PENGUIN to propose new directions in the molecular characterization of chromatin interactions as well as in the definition of potential targets for molecular screening towards disease treatment.

## Results

### The PENGUIN framework

PENGUIN builds EPINs by leveraging multiple sources of information. Specifically, it integrates diverse datasets, including (1) high-resolution chromatin interaction maps that capture active promoter-enhancer interactions, highlighting the dynamic nature of gene regulation; (2) tissue-specific physical nuclear protein-protein interactions (PPIs), enabling the exploration of the intricate molecular associations within the nucleus; (3) curated binding motifs of protein-DNA interactions, providing insights into the specific interactions between proteins and DNA and (4) gene expression levels, identifying active elements with the interaction networks (**Figure 1**). With this comprehensive approach, PENGUIN reconstructs EPINs by clustering enhancers that interact with the same promoter based on PPIs. Each EPIN consists of three distinct types of nodes: promoter-bound nodes, encompassing proteins with DNA binding motifs present in the promoter region; enhancer-bound nodes, comprising proteins with DNA binding motifs in the enhancer sequences; and intermediate nodes, representing proteins that interact with either the promoter-bound or enhancer-bound nodes but lack direct DNA binding motifs on the promoter or enhancers. By integrating these diverse nodes, PENGUIN provides a holistic view of the intricate molecular landscape within EPINs. This approach enables the exploration of the interplay between DNA-binding proteins, enhancers, and intermediate proteins, shedding light on the regulatory mechanisms that shape gene expression and ultimately influence cellular functions.

**Figure 1.**
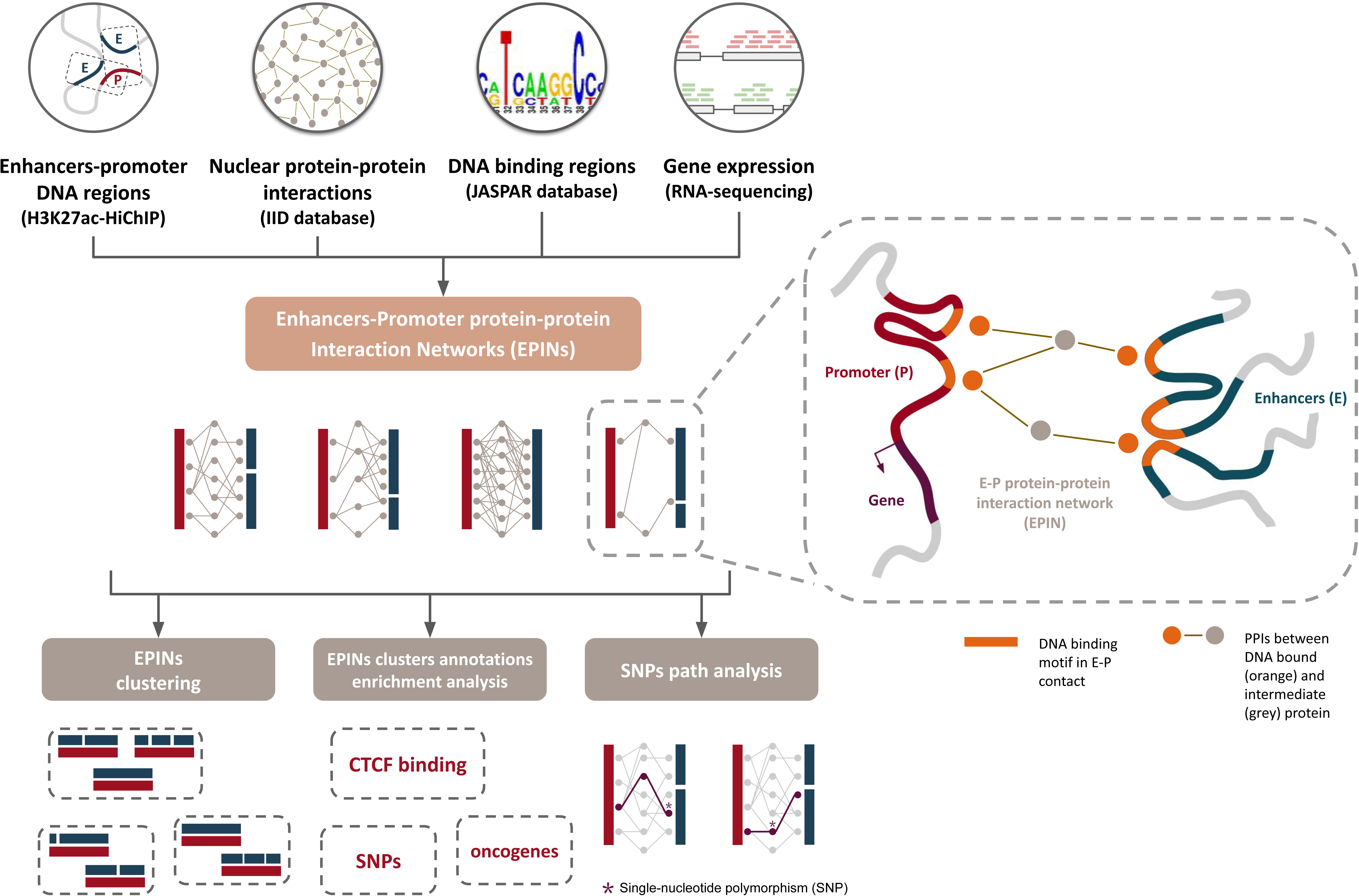
General overview of the PENGUIN workflow and downstream analyses. PENGUIN input consists of HiChIP data (in this work, H3K27ac in LNCaP or LHSAR cell lines), tissue-specific nuclear protein-protein interactions, PPIs (in this work, cancer and normal prostate PPIs from IID database), curated DNA-binding motifs (in this work, motifs from JASPAR database), and gene expression profiles (in this work, RNA-sequencing data in LNCaP or LHSAR cell line). PENGUIN output consists of Enhancer-Promoter protein-protein Interaction Networks (EPINs). Downstream analyses are designed to address specific questions related to prostate cancer (PrCa), namely the identification of clusters of promoters based on EPIN similarity, their enrichment in distinct annotations (CTCF binding from ChIP-seq peaks, PrCa-associated SNPs, and PrCa oncogenes), and finally the formulation of mechanistic hypothesis based on SNPs path analysis. In the inset, we report a schematic representation of an enhancer-promoter protein-protein interaction network (EPIN) reconstructed with PENGUIN for a given E-P contact detected by H3K27ac-HiChIP. Promoter and enhancer DNA binding motifs found in HiChIP regions after enhancer prioritization and the corresponding bound proteins are indicated in orange; their physical interactions with other factors of the EPIN (in gray) are represented as gray lines.

### PENGUIN identifies PrCa clusters of protein interactions based on chromatin contacts

We leveraged 24,547 E-P contacts (30,416 after refinement and prioritization, **Methods; Figure S1**) identified using H3K27ac-HiChIP data in LNCaP, 810 binding motifs from 639 DNA-binding proteins, and 31,944 prostate-specific, experimentally validated, physical and nuclear PPIs (filtering out proteins from unexpressed genes, **Methods; Figure S2**) to construct 4,314 EPINs using the PENGUIN clustering approach outlined in **Figure 2** (**Methods**). Each EPIN is centered around one promoter that we found to be contacted by a median of 4 enhancers, with a maximum of 93 enhancers for the promoter of the gene *CRNDE* (**Table S1**). Altogether, the 4,314 EPINs contain a total of 8,215 interactions (edges) among a total of 885 proteins (nodes) that are expressed in LNCaP (**Methods**). A mean of 36% proteins found in these EPINs are encoded by differentially expressed genes in LNCaP versus LHSAR (**Methods** and **Table S1**).

**Figure 2.**
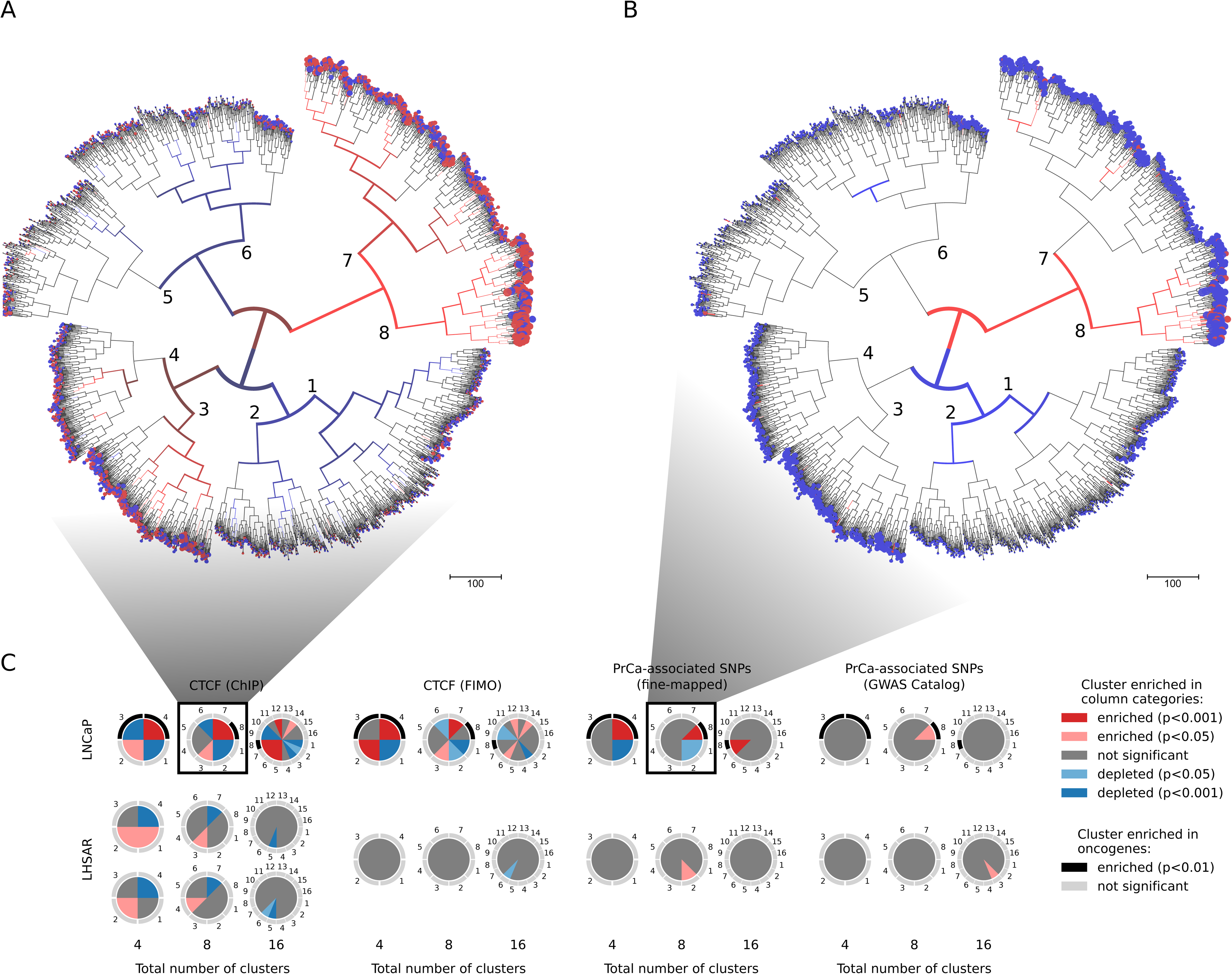
Clustering of the promoters originating the PENGUIN reconstructed EPINs. Clustering is based on edge composition of the EPINs. Leaf radius is proportional to network size. Color code (two-sided Fisher’s exact test): red, enriched; blue, depleted; The figure is generated using ETE3 ^68^. **(A**) Enrichment of PrCa SNPs in enhancers. We identified one PrCa SNP enriched cluster (GWAS+; cluster 8), and multiple PrCa SNP depleted (GWAS-; clusters 1, 2) and neutral (GWAS=; clusters 3, 4, 5, 6, 7) clusters. (**B**) Enrichment of CTCF ChIP-seq binding sites. We identified multiple CTCF enriched (CTCF+; clusters 3, 7, 8), depleted (CTCF-; clusters 1, 2, 6) and neutral (CTCF=; clusters 4, 5) clusters. (**C**) Clustering analysis on LNCaP (Top) and LHSAR (bottom) reconstructed EPINs. Pie-charts represent clustering results for a distinct total number of clusters used to partition the hierarchical clustering tree (4, 8, 16). Numbered pie-slices represent the different clusters and their color gradients encode the significance of enrichment (shades of red), depletion (shades of blue) or neutral (gray) of the overlap with distinct annotations (ChIP-Seq CTCF peaks, predicted CTCF binding sites by FIMO, PrCa-associated SNPs from fine-mapping and GWAS). Clusters significantly enriched with previously known oncogenes are annotated with black arcs. All enrichments have been estimated using two-sided Fisher’s exact test.

Overall, 751 out of the 885 proteins represent intermediate nodes, with 127 of them acting both as intermediate and as DNA-bound nodes in different EPINs (**Table S2**). 261 unique DNA-binding proteins have predicted binding sites in at least one of the anchors of enhancers and promoters. A mean of 32.8 (s.d. 11.5) distinct DBPs were identified per promoter anchor with SP1, EGR1, SP2 being the most represented; and a mean of 24.8 (s.d. 7.69) were predicted per enhancer anchor with SP1, IRF1 and TFAP2A being the most represented.

A mean of 1.43 (normalized) promoters (0.88 s.d.) are shared among enhancers, with a maximum of 15 promoters for the same enhancer. To identify communalities and differences among the 4,314 EPINs in LNCaP, we performed an unsupervised, hierarchical clustering based on edge composition (Ward’s linkage method, **Methods**). Using this approach, we identified 8 clusters of promoters with specific networks (**Table S1**,**Table S3, Figure 2 and Figure S3** and **Figure S4**).The decision to divide the hierarchical tree into 8 clusters was based on the analysis of cluster characteristics, achieved by varying the number of clusters (**Figure 2C**).

### Characterization of PrCa clusters identified by PENGUIN

We characterized the 8 clusters using PrCa specific annotations. We used the previously described 95% credible set of SNPs (henceforth referred to as PrCa SNPs) across 137 PrCa-associated regions fine-mapped from the largest publicly available GWAS summary statistics (N=79,148 cases and 61,106 controls ^19^). By comparing each cluster with all other clusters, we found a significant enrichment of PrCa SNPs in one specific cluster (cluster 8 or *GWAS+ cluster*; two-sided Fisher’s exact test, **Methods**). Interestingly this enrichment is exclusively due to SNPs in enhancers (**Table 1**). Our results show that E-P interactions containing PrCa SNPs are clustered together (red branches in **Figure 2A**) indicating that they have similar characteristics in the way their PPI networks are wired. We found that most pairwise interactions (67.5%, or 5,550 out of 8,215 edges) are found in all clusters but establishing different topologies.

**Table 1.** Enrichment of PrCa SNPs in cluster 8 (GWAS+) when considering SNPs overlapping enhancers, promoters, either or both.

| PrCa SNPs overlaps | Odds Ratio (OR) | p-value |
| --- | --- | --- |
| Only enhancers | 11.329 | 1.80e-12 |
| Only promoters | 1.139 | 0.6 |
| Either enhancers or promoter | 8.551 | 2.68e-11 |
| Both enhancers and promoter | 0 | 1 |

We identified the protein interactions that are enriched in each cluster and estimated the significance of overrepresentation of each edge in a cluster compared to all others (**Methods**). *GWAS+ cluster* (cluster 8 in **Figure 2; Figure S5**) exhibits the lowest number of promoters and distinctive network characteristics (**Table S3A, Figure S3**). Nonetheless, per promoter, it displays the largest number of edges (p-value < 1e-16) and intermediate nodes (p-value < 1e-16), in line with its greater number of enhancers per promoter (p-value < 1e-16), see **Figure S4**.

We then assessed whether PENGUIN clustering was influenced by super-enhancer-like regions sharing target promoters in given clusters. Although the distribution of enhancers per hotspots is similar among our 8 clusters (**Figure S4G**), the *GWAS+ cluster* has fewer single enhancers (enhancer at more than 15 kb from any other enhancer). The average number of promoters targeted by each hotspot for all our 3,752 defined enhancer hotspots was 1.83 promoters targeted per hotspot. When measured considering only the promoters in given EPIN clusters, the values were: 1.29 for cluster 1, 1.28 for cluster 2, 1.25 for cluster 3, 1.24 for cluster 4, 1.22 for cluster 5, 1.21 for cluster 6, 1.34 for cluster 7 and 1.27 for cluster 8. In this case, values were very similar between EPIN clusters.

Moreover, the EPINs of the GWAS+ cluster have the lowest values of node-level centrality measures, namely betweenness and degree (**Figure S3**). The degree of a node measures the amount of connections it has, while the betweenness centrality measures the amounts of shortest paths that pass through it. Low values of betweenness and degree indicate a lower amount of connections among different nodes of the network. Betweenness and degree are significantly different across clusters (Kruskall-Wallis test p-value < 1e-16), but not with respect to the ensemble of all EPINs, which indicates that, despite the high number of shared pairwise interactions (67.5% of edges), the wiring of the cluster-specific EPINs are distinctive.

Since CTCF is a major actor in the formation and maintenance of transcriptionally productive E-P interactions ^20, 21^, we tested the clusters identified by PENGUIN for enrichment in CTCF binding. For this analysis we used CTCF ChIP-seq peaks, from the same cell line (LNCaP), from the ENCODE project instead of predictions based on DNA-binding motifs (**Methods**). We found that the enriched interactions with CTCF peaks, that we call *CTCF+,* cluster together (red branches in **Figure 2C, Figure S5**), suggesting that the presence of CTCF in chromatin interactions results in the formation of characteristic PPI networks between the promoter and its enhancers.

CTCF+ clusters overlap the GWAS+ cluster (**Figure 2C, Figure S5**), suggesting that CTCF-mediated interactions could be more functionally relevant to PrCa. In particular, GWAS+ cluster (representing 6% of the total number of promoters considered) is the only one presenting the unique and significant enrichment in CTCF binding, PrCa SNPs, and oncogenes coincidentally (**Table 2, Table S3, Figures S5**). This cluster is enriched in the *Hippo signaling pathway* (KEGG:04390) (Bonferroni-corrected p-value=1.56e-3), *WNT Signaling Pathway* (KEGG:04310) (Bonferroni-corrected p-value=9.57e-3) and *Pathways in cancer* (KEGG:05200) with genes such as *BCL2L1, MYC, FOS* (Bonferroni-corrected p-value = 0.047) (**Methods, Table S5**). Interestingly GWAS+ cluster, or any other cluster, did not significantly stand out in terms of overall expression level (**Figure S2**) or, notably, in terms of fraction of differentially expressed genes (**Figure S2**).

**Table 2.** Enrichment of PrCa SNPs, CTCF ChIP-seq binding sites (“CTCF” in the header), and other PrCa annotations (oncogene promoters and PrCa SNPs from GWAS Catalog) across the eight clusters identified by PENGUIN. Cluster 8 is enriched in CTCF binding, PrCa SNPs, and oncogenes. Symbols code: +, enriched; -, depleted; =, neutral. OR: Fisher’s exact test Odds Ratio.

| Cluster | Number of genes | CTCF | OR CTCF | P-value CTCF | PrCa SNPs | OR PrCa SNPs | P-value PrCa SNPs | Number of oncogene promoters | OR oncogenes | P-value oncogenes |
| --- | --- | --- | --- | --- | --- | --- | --- | --- | --- | --- |
| 1 | 825 | - | 0.617 | 1.91e-9 | - | 0.28 | 2.46e-2 | 8 | 1.17 | 0.67 |
| 2 | 399 | - | 0.613 | 9.65e-6 | - | 0.00 | 2.00e-2 | 5 | 1.54 | 0.38 |
| 3 | 544 | + | 1.348 | 1.35e-3 | = | 0.80 | 8.27e-1 | 2 | 0.39 | 0.31 |
| 4 | 491 | = | 1.084 | 4.09e-1 | = | 0.51 | 3.60e-1 | 4 | 0.94 | 1.00 |
| 5 | 465 | = | 0.841 | 9.12e-2 | = | 0.75 | 8.14e-1 | 1 | 0.23 | 0.17 |
| 6 | 641 | - | 0.664 | 4.24e-6 | = | 0.38 | 1.03e-1 | 1 | 0.16 | 0.03 |
| 7 | 676 | + | 1.655 | 2.12e-9 | = | 1.42 | 3.18e-1 | 5 | 0.84 | 1.00 |
| 8 | 273 | + | 3.287 | 3.64e-20 | + | 11.33 | 1.80e-12 | 11 | 6.48 | 1.04e-5 |

To explore the potential connection between our clustering approach and the presence of trans-eQTLs, we used the trans-eQTLs reported from the largest eQTL study available (large-scale meta-analysis in up to 31,684 blood samples from 37 eQTLGen Consortium cohorts in Whole Blood, ^22^) and defined a region an ‘eQTL hotspot’ when associated to more than 3 genes (**Methods**). We observed an enrichment of eQTL hotspots across all clusters (**Figure 5SE**, empirical p-value <0.0001), but not specifically for cluster PrCA GWAS+ (**Figure 5SF**).

In conclusion, PENGUIN enabled the identification of a cluster of E-P contacts whose EPINs are uniquely enriched in PrCa SNPs, ChIP-seq CTCF peaks, and oncogenes (a.k.a. GWAS+ cluster or cluster 8, **Figure 2** and **Table 2**). It should be emphasized that our findings demonstrate consistent results also when employing PrCa-associated SNPs from the GWAS catalog, in which case we also identified cluster 8 as significantly enriched (**Methods, Table S6**).

### Baseline comparisons and assessment of PENGUIN specificity

Among the 273 promoters belonging to the identified GWAS+ cluster (cluster 8 in **Figure 2A**), 11 belong to known oncogenes, *FOXA1, ZFHX3, CDKN1B, KDM6A, BRCA2, CDH1, CCND1, NKX3-1, BAG4, MYC, GATA2* (**Methods**). We compared enrichment of PrCa functional annotations in the reconstructed networks with and without inclusion of intermediate proteins. Including intermediate proteins allows increasing the number of retrieved PrCa-related oncogenes in GWAS+ cluster from 6 to 11 and increasing significance of enrichment indicating improved specificity (**Table S4**). We then compared our results with the simple overlap of the genomic regions of E-P contacts and known oncogene promoters [see **Table S1**, which also reports on the overlaps of E-P contacts with CTCF peaks (in both enhancers and promoters, see **Methods**), and PrCa SNPs (in enhancers)]. In this scenario, only 30 promoters (12 overlapping the GWAS+ cluster) would be identified that overlap both PrCa SNPs and CTCF peaks. Of these, just 3 are promoters of known oncogenes (and only one, *ZMYM3*, is not in the GWAS+ cluster).

To explore the cell and disease-specificity of our results we applied PENGUIN on LHSAR, a benign prostate epithelial cell line. We performed H3K27Ac HiChIP experimental data and applied the PPI clustering procedure to explore functional relationships within the clusters. We then proceeded to apply PENGUIN to identify clusters of EPINs based on their edges (**Methods**). As the selection of an exact number of clusters in a given tree could be considered an important variable in our analysis, we examined various cluster numbers (4, 8, 16). We investigated the presence of cluster enrichment in GWAS and CTCF (**Table S3B**). Our analysis did not reveal any cluster enrichment in GWAS and CTCF within the benign prostate control LHSAR. Moreover, we did not observe a significant increase in the number of identified oncogenes in LHSAR (**Figure 2B**). These results lead us to conclude that PENGUIN, along with the integration of intermediate PPI networks, significantly enhances the identification of candidate PrCa-related SNPs affecting key elements in chromatin architecture.

Despite the high similarity in PPIs between LHSAR and LNCaP cells (Jaccard index of 0.85), their clustering based on H3K27Ac HiChIP data revealed distinct EPINs (**Figure 2B**). This finding highlights the sensitivity of our method in capturing subtle differences within EPINs. To further validate this, we conducted additional statistical analyses on PPIs across different cancer cell types. By examining the overlap between PPI networks, we discovered significant variations that were highly specific to each cell type (**Figure S6**). This observation not only reinforces the reliability of the differences found in LHSAR and LNCaP cells but also suggests that our results can be expected in other cellular contexts provided the required H3K27ac-HiChIP information, which is currently unavailable in most cases.

To further investigate the significance of intermediate PPI networks, we conducted clustering analysis exclusively based on HiChIP interactions. Specifically, we utilized the list of enhancer IDs, denoted by their genomic coordinates, within each EPIN (**Figure S7**). Our findings unequivocally demonstrate that the exclusion of intermediate PPI networks substantially diminishes the number of identified oncogenes. This outcome strongly suggests that the information conveyed by the PPI network plays a crucial role in the classification of EPINs and their correlation with phenotypic traits.

### Involvement of E-P protein interactomes in tumor-related functional processes

We analyzed the functional enrichment of the set of 885 proteins composing the universe of nodes used in the EPINs of LNCaP. 43 out of these 885 proteins are encoded by one of the 122 known PrCa oncogenes (32 intermediates, 7 DBPs among which MGA, ETV4, ETV1, GATA2, ETV3, ERF, NKX3-1, and 4 of both types among which TP53, MYC, FOXA1, AR; see **Methods** and **Table S2**). In total, 11 out of 885 have been targeted by PrCa-specific drugs (source: DrugBank; protein targets: ESR2, ESRRA, AR, PARP1, NFKB2, NFKB1, NCOA2, NCOA1, AKT1, TOP2A, TOP2B; drugs: Estramustine, Genistein, Flutamide, Nilutamide, Bicalutamide, Enzalutamide, Olaparib, Custirsen, Amonafide); and 190 out of 885 are targets of non-prostate drugs indicating the possibility of re-purposing.

Considering the genes encoding for 477 out of 751 intermediate proteins with annotations for KEGG pathways retrieved using g:Profiler ^23^, 41 were annotated in the prostate cancer pathway (KEGG:05215) (adjusted p-value = 3.62E-24), which annotates a total of 97 genes (**Methods** and **Table S7**). We next studied specific protein enrichments in the nodes of the EPINs of each identified cluster (**Table S8**). Although intermediates are ubiquitous and generally shared among all clusters, we could identify 22 significantly specific proteins enriched in the GWAS+ cluster (**Methods**). Functional enrichment analysis of these 22 proteins revealed significant relationships with tumorigenic processes (**Table S9**). KEGG *Prostate cancer pathway* (KEGG:05215) appears highly enriched (adjusted p-value = 1.27e-2) together with other pathways related to tumors such as *Colorectal cancer* (KEGG:05210, adjusted p-value = 3.20e-5) *Pancreatic cancer* (KEGG:05212, adjusted p-value = 9.54e-4) and *Breast cancer* (KEGG:05224, adjusted p-value = 7.06e-4). KEGG pathway KEGG:04919 (*Thyroid hormone signaling pathway*) is an additional highly enriched pathway (adjusted p-value = 2.57e-4). Thyroid hormones have been previously described as modulators of prostate cancer risk ^24–27^. Pathway KEGG:05200 (called *Pathways in cancer*) appears as the fourth most enriched KEGG concept (adjusted p-value= 3.63e-4). Other classical tumorigenic pathways, such as *Wnt signaling pathway* (KEGG:04310, adjusted p-value = 1.27e-2) and *TGF-beta signaling pathway* (KEGG:04350, adjusted p-value = 8.21e-4) appear to be enriched. In this regard, recent studies analyzed the involvement of Wnt signaling in the proliferation of prostate cancer cells ^28, 29^, as well as the involvement and TGF-beta signaling ^30, 31^.

Furthermore, we examined the functional enrichment of significantly central proteins across all other clusters. This analysis was conducted to facilitate functional comparisons across different clusters (**Methods ‘**Functional gene set enrichment analysis’). This analysis revealed no enrichments for clusters 1, 2, 4, 5, and 6 (cluster 5 does not have significantly central proteins). This observation can be attributed to the higher number of central proteins in these clusters (365 in cluster 1, 283 in cluster 2, and 318 in cluster 6) compared to the other clusters (3 in cluster 3, 7 in cluster 7, and 22 in cluster 8). Despite having a similar number of significantly central proteins to cluster 8 (30 proteins), cluster 4 does not show any enrichment.

Moreover, of the clusters presenting enrichments (i.e., clusters 3 and 7), only cluster 7 presents enrichments related to those observed in cluster 8 (for example, KEGG *prostate cancer pathway* is enriched, adjusted p-value = 2.041e-2; **Figure S8**). As commented, cluster 7 presents only 7 significantly central intermediate proteins (CREBBP, CTNNB1, GSK3B, KAT5, MAPK1, PIN1, SMAD2), out of which, 6 overlap with those significantly central in cluster 8 (only PIN1 is absent).

### SNPs path analysis in the E-P protein interactomes

Next, we sought to perform an analysis of the SNPs found along the paths within each EPIN (**Methods**). In this analysis, a path in a network is a sequence of edges joining a sequence of nodes connecting the promoter and the enhancers of an EPIN (**Figure 3A**). We distinguish between two possible scenarios based on the location of the SNPs within the paths: (1) PrCa SNPs fall in the DNA binding motifs found in enhancers, indicating a possible dysregulation of TFs binding and activity (**Figure 3B**); (2) PrCa SNPs in the genomic regions of the genes that encode for the intermediate nodes of the EPINs, indicating a possible alteration of the PPIs (**Figure 3C**). The first analysis aims to identify the location of enhancers that could be targeted by genetic perturbation techniques such as CRISPR/Cas9. The second analysis aims to identify the proteins that are potentially affected by mutations so as to enhance our understanding of prostate cancer biology. Overall, we characterized all PrCa SNPs falling within any path that connects enhancers to a promoter (rs4962419 was found in both scenarios analyzed). In the following, we discuss the two scenarios and report on the *MYC, CASC11 and GATA2* promoters as illustrative examples.

**Figure 3.**
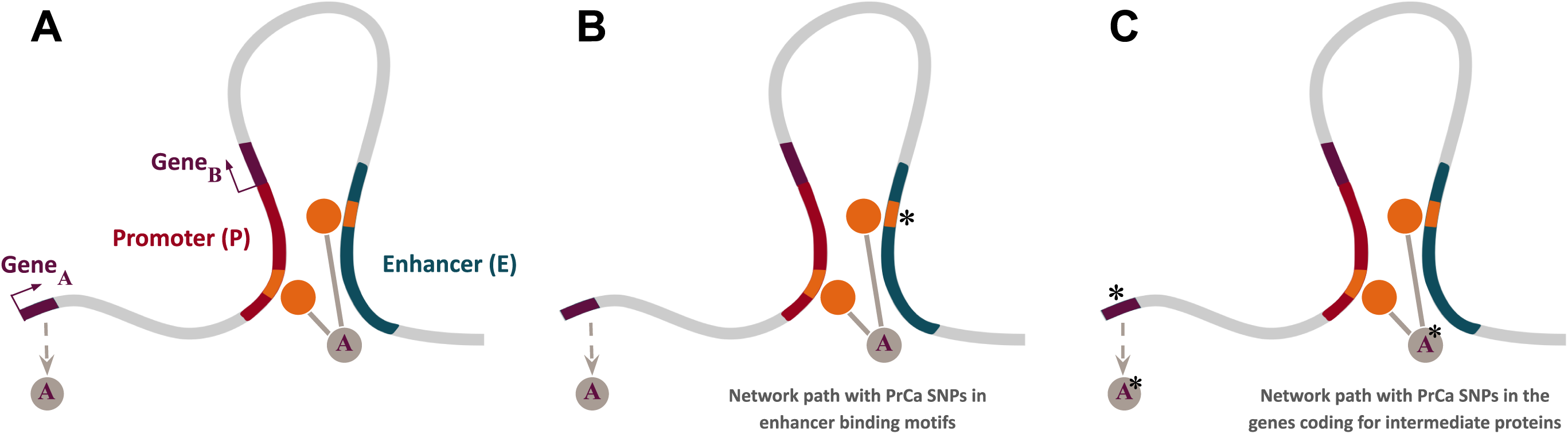
Schematic representation of different types of network paths found in the EPINs reconstructed by PENGUIN. In general, a network path is defined by an intermediate protein (gray circle), encoded by a gene (dark red line; Gene_i_), that interacts with DBPs (orange circles) with binding motifs (orange lines) on the enhancer (green line) and the promoter (red line) of another gene (dark red line; Gene_j_) (**A**). If a PrCa SNP (asterisk) falls in the enhancer binding motif, the interaction between the DBP and the enhancer may be disrupted and possibly its interactions (**B**). If a PrCa SNP (asterisk) falls in the gene that encodes for the intermediate protein, the gene product could be affected and possibly its interactions **(C)**. Colors are consistent with Figure 1.

### Network paths with PrCa SNPs in enhancer binding motifs

We sought to detect SNPs located in the DNA binding motifs found in the enhancers of the EPINs. Based on previous evidence ^32, 33^, our hypothesis is that SNPs in enhancers could disrupt the binding of proteins such as TFs having an impact on their interactome.

In **Table S10** we list the 36 PrCa SNPs falling within 60 DBP motifs in enhancer regions linking 34 different promoters whose EPINs include 5,184 edges. Among these, we identified 17 PrCa SNPs falling within 16 EPINs (1,894 edges) belonging to the *GWAS+ cluster* that had at least one PrCa SNP in their enhancers. Several of these EPINs have promoters of differentially expressed genes (such as *DLL1, STOM and SEC11C* in the GEPIA tumor/normal dataset; *ID2, RPS27, SEC11C, CASZ1, CRTC2, C5 and STOM* in the LNCaP/LHSAR dataset; see **Methods, Differential Gene Expression**).

To establish the biological significance of the identified SNPs, we leveraged data from previous pooled genome-wide CRISPR/Cas9 knockout and RNAi screens conducted in prostate cancer LNCaP cells, available in the DepMap database (https://depmap.org/, DepMap ID: ACH-000977). These screens provide essentiality scores, which quantify the relevance of specific gene networks to the proliferation of LNCaP cells. In our analysis, we retrieved essentiality scores for genes in prostate tissue from DepMap and compared three distinct gene sets: (1) the genes (EPIN promoters) prioritized in **Table S10**, (2) all genes (EPIN promoters) included in our analysis, and (3) all genes available in the DepMap database. Remarkably, we observed significant differences in the essentiality scores (Z-scores) among these sets, with lower Z-scores indicating a higher degree of gene essentiality (**Figure 4A**). This analysis aligns with the RNAi findings, demonstrating a significant decrease in essential scores for genes containing the SNPs listed in **Table S10 (Figure 4B)**. Furthermore, the GSEA analysis unveiled a noteworthy enrichment (p-value = 0.0017) for these EPIN promoters that harbor intermediate nodes with SNPs at their genomic location (as indicated in the supplementary **Table S10**) (**Figure 4C**). Among the top essential genes, the CRISPR/Cas9 and RNAi screens prioritize the following ones : GATA2-AS1, CASZ1, MYC, KRT8, GTPBP4-AS1, MFN2, CTBP2, and ID2.

**Figure 4.**
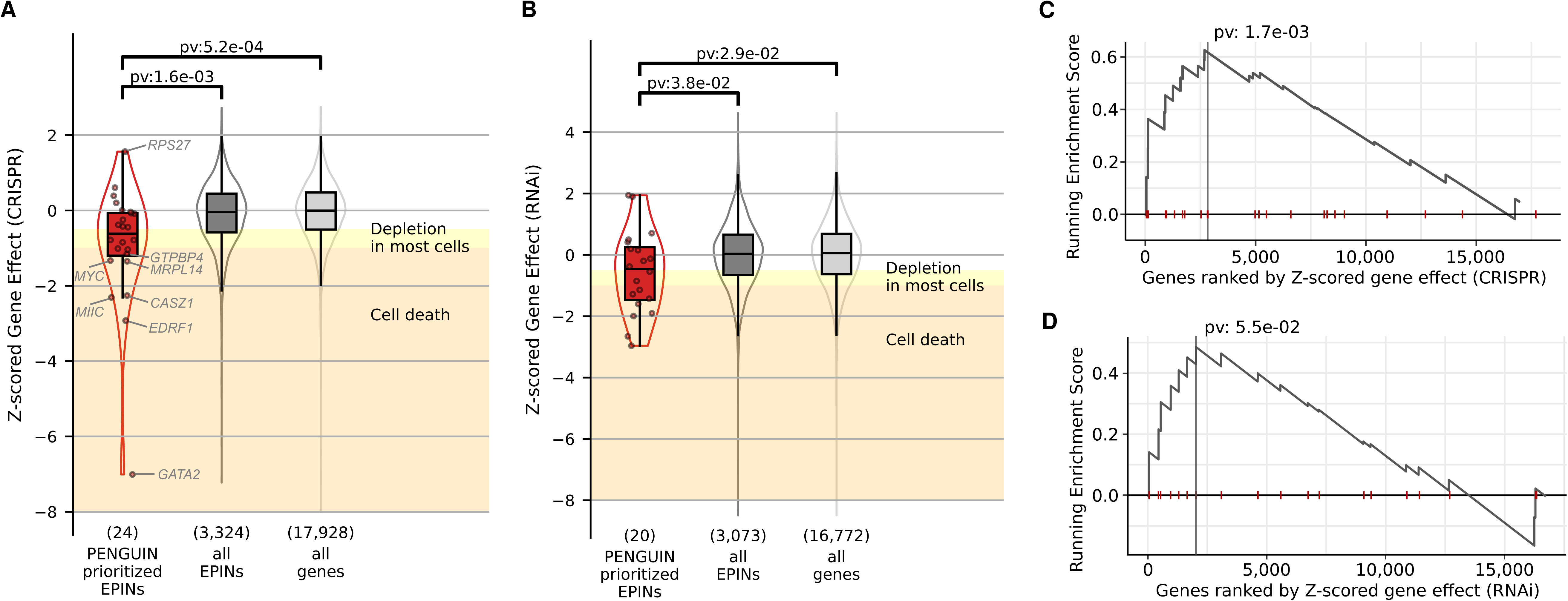
Validation of SNPs prioritized by PENGUIN. CRISPR/Cas9 knockout and RNAi screens provide Z-scores to quantify the relevance of a specific gene network to proliferation of LNCaP cells. **(A)** CRISPR/Cas9 knockout analysis indicates that intermediate SNPs prioritized by PENGUIN occur in genes essential for LNCaP (significance calculated with Mann-Whitney test). Genes with the strongest effect are displayed. **(B)** RNAi analysis shows milder but significantly consistent results with CRISPR/Cas9 knockout. **(C)** Gene Set Enrichment Analysis (GSEA) indicates that SNPs prioritized by PENGUIN occur in the most essential genes identified by CRISPR/Cas9 knockouts. (**D**) GSEA indicates that SNPs prioritized by PENGUIN occur in the most essential ones based on the RNAi screen. for C and D, the statistical significance of the enrichment of a gene set within the ranked gene list is reported.

Finally, at the level of intermediate proteins, we also found some encoded by genes reported to be differentially expressed. We observed that the mean proportion of intermediates that are differentially expressed is on average 40% (**Figure S4**). We tested whether promoters belonging to the GWAS+ cluster were significantly enriched for intermediate protein encoding for differentially expressed genes (**Methods**). Among the 16 EPINs belonging to the GWAS+ cluster that had at least one PrCa SNP in their enhancers, 11 contain expression data to study potential direct effects of the SNPs. In this subset we found 4 EPINs differentially expressed in promoters (3 also differentially expressed in intermediates: *CASZ1, ID2, SEC11C*), and 4 EPINs only differentially expressed in intermediates: *MIIP, MRPL14, MYC, TMEM63B* (**Table S1**). The differential expression of intermediates makes it easier to identify interesting and potentially novel cases. For instance, MYC is not differentially expressed but it has differentially expressed intermediates.

### Network paths with PrCa SNPs in the genes coding for EPIN nodes

In this analysis, we identify EPINs with PrCa SNPs falling within genes that encode either for intermediate or anchor bound nodes (**Table S11**), indicating a potential alteration of PPIs involved in E-P contacts. We found that the GWAS+ cluster has the highest proportion of PrCa SNPs in these nodes compared to all other clusters (mean = 53.2, SE = 18.0, p-value <= 0.01, **Table S12**). The EPINs of *STK40* and *GATA2* promoters in GWAS+ cluster display the highest fraction of EPIN nodes with PrCa SNPs in their corresponding genes encoding them (**Table S1**).

We use the SNP paths to link 172 PrCa SNPs falling within the gene bodies of 26 genes of which 7 are known oncogenes (*MAP2K1, CHD3, AR, SETDB1, ATM, CDKN1B, USP28*). We identify edges that are most enriched in our GWAS+ cluster which could be pointing to essential links between the gene encoding for the node and containing a PrCa predisposing SNP at a particular EPIN. For example, we identify the link between *MDM4* containing SNP rs35946963 (PrCa p-value 1e-24) and TP53 ^34^ and between *KDM2A* containing SNP rs12790261 (PrCa p-value 1e-7) and BCL6 ^35^ and *ARNT* continuing SNP rs139885151 (PrCa p-value 3e-13) and HIF1A ^36^.

We integrated information from pQTL associations between the 172 PrCa SNPs and protein levels (**Methods**). Two intermediate proteins (CREB3L4, MAP2K1) were associated with PrCa SNPs falling within the gene encoding for them (p-value of association with proteins were 7.75e-86 for CREB3L4 and 2.40e-5 for MAP2K1). We identified 3 out of 26 promoter EPINs (*TRIM26, MEIS1, POU2F2*) with suggestive evidence (p-value < 1e-5) of association between the PrCa SNP with the PENGUIN-linked promoter EPIN, pointing to the cancer promoting mechanistic action of these variants: gene with SNPs in *POU2F2* linked to the EPIN promoter of gene *PHGDH* (SNP with lowest p-value rs113631324 = 3.80e-8); gene with SNPs in *TRIM26* and EPIN promoter of gene *RRM2* (SNP with lowest p-value rs2517606 = 2.69e-7); gene with SNPs in *MEIS1* and EPIN promoter of gene *STOM* (SNP with lowest p-value rs116172829 = 8.19e-6).

We note that, unlike SNPs in enhancers, whose effect can be directly assessed by CRISPR/Cas9 or RNAi assays, the impact of SNPs on intermediate nodes is more complicated to estimate due to their shared involvement in multiple gene networks. In fact, it is worth mentioning that among the 885 proteins identified by PENGUIN, 751 serve as intermediate nodes (section **PENGUIN identifies PrCa clusters of protein interactions based on chromatin contacts**). This overlapping functionality further complicates the prediction of SNP effects on these intermediate nodes.

### Examples: SNPs path analysis of *MYC*, *CASC11* and *GATA2* promoters

From HiChIP data, the *MYC* promoter (chr8:128747814-128748813) is in contact with 73 enhancer regions among which one holds the SNP rs10090154 (p-value of association with PrCa = 1.4e-188). This SNP is located in the binding motif of the transcription factor FOXA1 on the *MYC* EPIN enhancer. The integration of PrCa SNPs information highlights paths in the EPIN of *MYC* that are particularly compelling in the context of the disease (red line in **Figure 5; Figure S9**). The promoter region of *MYC* binds 8 proteins TFAP2C, KLF5, RBPJ, SP1, ZBTB14, ATF6, ZBTB7A, PRDM1 and contains 17 protein interactors (dots in **Figure 4**) that might be affected by the possible disruption of its binding motif, namely, HMGA1, RCC1, TFAP4, NFIC, PBX1, HOXB9, NFIX, NACC1, RARA, PIAS1, RPA2, H2AFY, RECQL, SATB2, CREB1, AR. The gene encoding for *FOXA1* is differentially expressed, along with others of its interactors (**Table S10**; **Methods**). Interestingly, 24 PrCa SNPs fall within the genomic region of *AR* (marked by an asterisk next to the gene name), all with p-values of association with PrCa below 1e-11 (**Table S11**). AR is targeted by several drugs used in the treatment of prostatic neoplasms, such as apalutamide, bicalutamide, diethylstilbestrol, enzalutamide, flutamide, and nilutamide (triangle in the **Figure 4A**, source: DrugBank). Notably, mutations in *FOXA1* enhancers were previously shown to alter TF bindings in primary prostate tumors ^33^. And, also in line with our observations, *FOXA1* enhancer region has been previously reported to be coupled to *MYC* ^37^ and has been shown to have a strong binding of AR ^38^.

**Figure 5.**
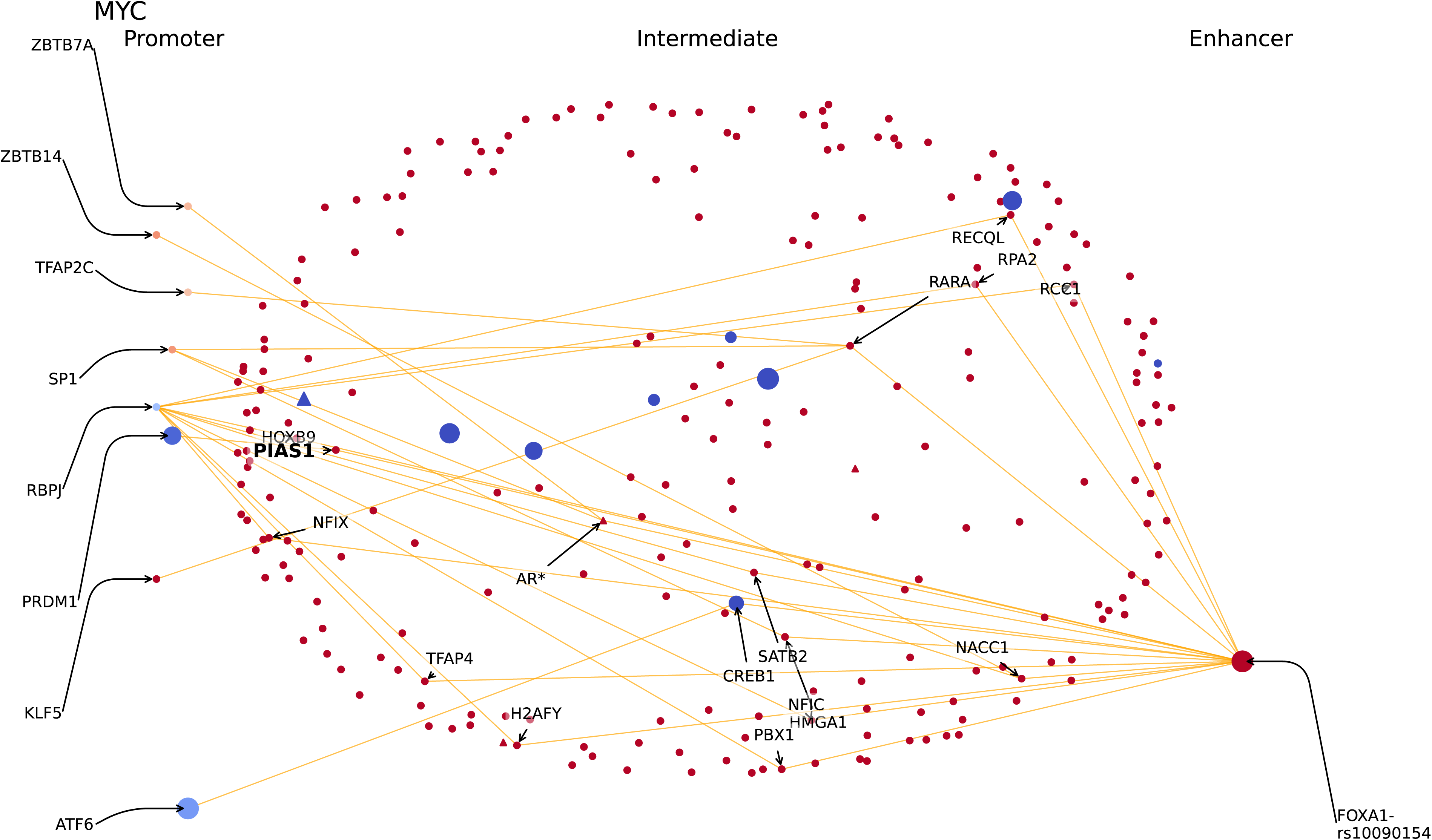
Reconstructed protein interactions between *MYC* promoter and its enhancers. DBPs with binding motifs on the promoter region are aligned on the left, while those with binding motifs on the enhancers are aligned on the right. In the middle, proteins that connect DBPs through a shortest path. Each dot represents a protein. Color, size and shape codes are explained in the Tutorial section of the PENGUIN web service at https://penguin.life.bsc.es/. In this figure, only the edges of network paths with PrCa SNPs in enhancer binding motif are represented (orange lines). Such PrCa SNPs are indicated beside the name of the enhancer-bound DBP (e.g. FOXA1-rs10090154); PrCa SNPs in intermediate proteins are indicated with an asterisk (e.g. AR); the proteins found to be enriched in the GWAS+ cluster are highlighted in bold (e.g. PIAS1); druggable proteins from DrugBank are indicated as triangles.

We report two additional examples, the EPINs for the promoters of *CASC11* (**Figure S10A**) and *GATA2* (**Figure S10B**). The EPIN of *CASC11* promoter is also affected by variant rs10090154, the same well known variant associated with risk of developing prostate carcinoma that we introduced with *MYC* EPIN ^39, 40^(**Table S10).** Interestingly, *CASC11* is known to enhance prostate cancer aggressiveness and is regulated by C-MYC ^41^, while being close to the *MYC* gene on chromosome 8. The promoter binds 6 proteins: TFAP2C, SP3, SP1, PKNOX1, NR2C2 and KLF5. Potentially affected protein interactors of the EPIN include: HMGA1, PIAS1, AR, RARA, and PBX1. GATA2 is an interesting case given its essentiality score from DepMap (Z-score=-7.01). Its EPIN presents up to 11 intermediates affected by PrCa related SNPs, namely TCF4, CTBP2, AR, ARNT, TCF7L2, CDKN2A, NEDD9, ANKRD17, MEIS1, MDM4 and CHD3. The role of GATA2 as mediator of AR signaling in AR-dependent prostate cancer, as well as its role as a potential target for treatment development ^42^ has been previously described, as silencing of the gene is known to affect other relevant genes such as *C-MYC* and *AURKA* (Chiang et al., 2014). Proteins bound to the promoter region include: ZBTB7A, ZBTB33, TCF3, SF1, NR2C2, KLF3, EGR1, E2F1 and CREB1, but most importantly, the EPIN presents AR bound to the enhancer region, which, as we pointed out with *MYC* EPIN, is the target of several PrCa treatments.

## Discussion

Here we introduced the PENGUIN approach that operates on the premise that the EPIN network structure connecting a promoter and its enhancers can serve as a distinctive signature associated with specific functional profiles and diseases. Our assumption is grounded in earlier research that has demonstrated the correlation between 3D loop topology and chromatin state or gene expression ^43^. We propose PENGUIN as a molecular approach to study variations in structural characteristics of chromatin loops, establishing a direct link to disease-related phenomena. By integrating the PPI network information, the method offers valuable insights into the underlying mechanisms driving these distinctive features and their relevance to disease progression.

Previous approaches have already incorporated PPI networks with GWAS hits to enhance their analysis. For instance, Ratnakumar et al. ^44^ identified proteins that exhibited an enrichment of PPIs with GWAS hits. In a recent study, Dey et al. ^45^ demonstrated the benefits of employing strategies that capture both distal and proximal gene regulation in prioritizing disease-related genes. In addition, alternative methods have amalgamated information from 3D chromatin interactions and GWAS SNPs to establish connections between intergenic SNPs and gene regulation in cancer contexts ^3, 46, 47^. These approaches have contributed to unraveling the relationship between genetic variations, chromatin organization, and disease. In contrast, our method takes a unique approach by being completely agnostic to the presence of SNPs. It combines information from PPI networks and enhancer-promoter interactions derived from H3K27ac-HiChIP data within a unified framework. This integrative approach allows us to leverage both the protein interaction landscape and the regulatory interactions between enhancers and promoters, leading to a comprehensive understanding of the molecular mechanisms underlying disease.

By utilizing PPI networks, we were able to reveal a distinct set of genes associated with PrCa that would have remained undiscovered using other methods. Notably, the intermediate nodes within this PPI network possess intrinsic properties that can be leveraged for the classification and characterization of E-P chromatin loops. Thus, our study demonstrates the capability of PENGUIN to group genes based on their involvement in PrCa, even in the absence of any prior information. This breakthrough opens up an uncharted avenue towards comprehending and identifying unsuspected biological markers in disease. In particular, the genes identified within the cluster exhibiting the highest enrichment in SNPs associated with PrCa (the GWAS+ cluster) can be considered promising candidate oncogenes or potential partners of oncogenes. It is conceivable that these genes may share "onco-enhancers," which are enhancers contributing to tumorigenic activity. For instance, PENGUIN can be used to identify *trans-*acting factors (e.g., interaction cascades of TFs and chromatin regulators) that could be targeted by drugs, or *cis-*acting factors (e.g., DBPs with binding motifs in regulatory elements) whose DNA binding affinity could be modified through knock-outs via CRISPR for therapeutic intervention. Moreover, unlike traditional TF enrichment analysis which detects general enrichments of particular proteins, PENGUIN can help identify the specific protein cascade potentially disrupted at enhancer loci for the disease under study. Overall, our findings highlight the potential of PENGUIN in unveiling previously unknown gene networks and provide valuable insights into the identification and characterization of biomarkers in various diseases, including PrCa.

To validate our findings, we have used cell-line specific datasets, androgen-sensitive human prostate adenocarcinoma cells (LNCaP) or a normal prostate epithelial cell-line (LHSAR). Each of the sources of information could be directly or indirectly related to the specific cell-lines used in this study: (1) H3K27ac-HiChiP in LNCaP and in LHSAR, (2) prostate-specific PPIs and (3) DNA binding motifs extracted from publicly-available datasets but filtered by our cell-type specific interacting 1 kb promoter-enhancer regions and (4) gene expression on cell-line for filtering PPI networks. The comparison of the results in cancer cell-line (LNCaP) to the results in a benign cell line (LHSAR) support our PrCa cell-specific findings. In LHSAR we found a significant association between the obtained clusters and the presence of CTCF, pointing towards the correct classification of EPINs into biologically relevant categories. But, this same clustering in the benign LHSAR cell-line did not reveal any association to PrCA, neither at the level of PrCa-SNPs, nor at the level of specific oncogenes. Future analyses could explore the use of clustering E-P loops with PENGUIN using other methods and sources for each of these layers. For example, we have used as input enhancer-promoter loops cell-specific H3K27Ac HiChIP experiments (strict calling of loops and prioritization), to maximize our true positives in the input data. The input for the PENGUIN clustering approach can also be constituted by enhancer-promoter links measured from other experimental methods aside from HiChIP or even using computational methods. We leave this for subsequent analyses.

In this work, we use a targeted approach and use the information on association of SNPs from fine-mapping as an annotation to our clusters. Specifically, we identify potential SNP paths from defined PrCa associated regions. SNP paths link genes in a network through a path that either starts from TF binding sites in enhancers or passes through proteins from the intermediate EPIN network that would have SNP in their gene bodies. This approach adds a new dimension in the contextualization of GWAS-associated SNPs using the EPIN looping realm.

It is important to mention our primary objective was to shed light on specific links that could be disrupted by PrCa-predisposing variants, such as CTCF bindings that connect promoters to their enhancers, or intermediate structural proteins that play a role in the E-P network. Further investigation is required to gain a comprehensive understanding of the biology and mechanisms underlying these crucial links. For this purpose, and to facilitate the exploration of SNP pathways associated with prostate cancer, we developed a user-friendly web interface accessible at https://penguin.life.bsc.es/. This platform serves as a tool for convenient investigation into the pathways influenced by SNPs in the context of prostate cancer. It is also intriguing to observe that, while PENGUIN successfully identifies clusters of EPINs significantly associated with PrCA, the gene expression analysis did not reveal any significant trends. At first glance, this observation may appear contradictory to our definitions of EPIN clusters and the core concept of EPIN itself. However, considering the evidence presented by our analysis, we believe that PENGUIN enables the detection of cancer associations with heightened sensitivity compared to traditional differential expression analyses. The ability of PENGUIN to capture intricate associations between EPINs and cancer surpasses the limitations of relying solely on gene expression changes, offering a more comprehensive understanding of the underlying molecular mechanisms involved in cancer development and progression.

Our analysis comes with some caveats to keep in mind. Firstly, we relied on data from the HiChIP technique for capturing enhancer-promoter (E-P) interactions, protein-DNA interactions from FIMO, and tissue-specific protein-protein interactions from the integrated interactions database (IID). The comprehensiveness of these datasets is inherently limited by the scope and constraints of the underlying databases and methodologies employed. Furthermore, our approach focuses on networks involving proteins with known edges, resulting in a consideration of only those proteins. Additionally, for the purpose of visualization, we have condensed the number of reported proteins and have presented only one intermediate protein (expanded one edge away). Moreover, it is worth mentioning that our study focuses on E-P interactions within a stable environment (LNCaP cells), representing a snapshot in time. While this field is still undergoing active research and further exploration, existing literature suggests that E-P interactions can exhibit minimal and quantitatively small changes in these conditions. Thus, while interpreting our findings, it is essential to consider the limitations of the utilized databases and methodologies, the specific protein selection, the condensed visualization approach, and the stable cellular context in which the E-P interactions were examined.

In conclusion, the PENGUIN approach employed in this study to investigate PrCa in LNCaP cells has the potential to be applied to the study of other human diseases, given the availability of similar data. This approach can be extended to explore different scenarios, such as different cell types or combinations of GWAS data, offering a promising avenue for future investigations. For instance, utilizing E-P dataset from another prostate cancer cell line would allow the identification of target genes regulated by enhancers from diverse cell types. These target genes can be prioritized using a genome-wide collection of disease-specific risk SNPs. The networks generated by PENGUIN provide a molecular understanding of the associations involved in cancer-related chromatin dynamics, making them well-suited for training advanced machine learning models like graph neural networks (GNNs). We propose potential intermediates in PrCa that engage in E-P networks within cancer cells and present opportunities for therapeutic intervention. High-throughput functional studies could validate the impact of genetic perturbations on thousands of enhancers simultaneously. As shown in our analysis, leveraging CRISPR-Cas9 technology would enable precise editing of specific genomic regions, facilitating targeted investigations and further elucidating the functional consequences of these genetic perturbations.

## Acknowledgments

The authors are grateful to José María Fernández González (Barcelona Supercomputing Center) for the crucial guidance with the PENGUIN web server. They also thank the Biola Javierre’s lab at the Josep Carreras Leukaemia Research Institute for the support, the ‘RNA initiative’ at IIT and all the members of Tartaglia’s lab at CRG, Sapienza and IIT.

## FUNDING

The research leading to these results has been supported by the European Research Council [RIBOMYLOME_309545 and ASTRA_855923], the H2020 projects [IASIS_727658 and INFORE_825080], and the project ONCOLOGICS (ERA Net Grant 779282, ERAPERMED2020-036; and Departament de Salut-Generalitat de Catalunya support, SLD040/20/000001). CG has received funding from the European Union’s Horizon 2020 research and innovation programme under the Marie Skłodowska-Curie grant agreement No 754490 – MINDED project. I.N.C. was supported by a grant for pre-doctoral contracts for the training of doctors (Project ID: SEV-2015-0493-18-2) (Grant ID: PRE2018-083662) from the Spanish Ministry for Science, Innovation and Universities.

## Methods

### Conformation capture and E-P interactions

We used Hi-C followed by chromatin immunoprecipitation (HiChIP) targeting H3K27Ac in LNCaP cells (androgen-sensitive prostatic carcinoma cell line) across 5 biological replicates including 1 billion reads as previously described ^11^ (Table 3). As a comparison, we also performed H3K27Ac HiChIP on LHSAR (Prostate epithelial cells overexpressing androgen receptor), across three replicates including 309 million reads. HiChIP, an efficient protein-mediated chromatin-conformation assay, was performed following the procedure described ^10^. The alignment, processing and loop calling from raw fastq files (paired-end data) was performed as previously described ^11^. Briefly, HiC-Pro ^48^ was used to map the HiCHiP trimmed reads and extract unique interactions; FitHiChIP ^49^ was used to identify significant interactions with a predefined set of peaks from H3K27ac ChIP-seq in LNCaP to refine accurate anchor ranges. We used q-value < 0.01 and a 5 kb resolution and considered only interactions between 5 kb and 3 Mb as previously described ^11^. In this analysis, we restricted to a stringent global background estimation to reduce as much as possible the number of false-positive interactions. The corresponding FitHiChIP specifications used were “IntType=3” (the peak-to-all) for the foreground, meaning at least one anchor to be in the H3K27 peak, and “UseP2PBackgrnd=1” (the peak-to-peak (stringent)) for the global background estimation of expected counts and contact probabilities for each genomic distance for learning the background and spline fitting. We identified 49,565 significant interactions (FitHiChIP, FDR<0.01) for LNCaP, and 12,053 for LHSAR.

**Table 3.** Genomic datasets used in the work. Data with no references were generated for this study. (*) Not from LHSAR but from human epithelial cells of the prostate.

|  | LNCaP | LHSAR |
| --- | --- | --- |
| HiChIP H3K27ac | 5 replicates<sup>11</sup> | 3 replicates |
| RNA-seq | 2 replicates<sup>50</sup> | 2 replicates<sup>50</sup> |
| ChIP-seq H3K27ac | 1 replicate<sup>11</sup> | 2 replicates |
| ChIP-seq CTCF | 2 replicates<sup>51</sup> | 2 replicates<sup>*51</sup> |

We categorized interactions by overlapping anchors with transcription start sites (TSS) and enhancers identified by H3K27ac ChIP-seq as previously described ^11^. Briefly, we first extended anchors by 5 kb on either side; we defined promoter regions around the TSS (+/- 500 bases) using RefSeq hg19 (see **Data Availability**); we defined enhancer regions using regions from H3K27ac ChipSeq in the same cell. Specifically, these were 49,638 and 53,561 enhancer regions, respectively from H3K27ac LNCaP in regular media (union of narrow and broad peaks) and from H3K27ac LHSAR. We note that the enhancer anchors at this stage of the analysis are of length 15 kb, due to 5 kb resolution of the HiChIP data analysis and additional 5 kb padding added to anchors on either side. We labeled the promoters and enhancer regions that overlap either right or left anchors, and considered E-P if only one anchor overlaps a promoter and the other an enhancer region. For LNCaP, out of the 49,565 significant interactions, we considered 18,151 E-P interactions. For LHSAR, out of the 12,052 significant interactions, we considered 5,435 E-P interactions. It is important to emphasize that our study relies solely on enhancers defined by our own HiChIP experiments, rather than relying on annotated enhancers or external definitions from ENCODE. We further prioritized E-P interactions to 1 kb regions and discarded from enhancers the 1 kb bins with fewer HiChIP interactions with the promoter (see *E-P HiChIP prioritization* section). We obtain 30,416 and 4,497 E-P interactions of 1 kb each for LNCaP and LHSAR respectively. The 15 kb original E-P interactions dataset contained a mean of 1.6 (1.3 s.d.) promoter anchors per enhancer anchor (after prioritization of enhancer anchor to 1 kb region, mean of 1.4 (0.9 s.d.) promoters per enhancer). There were 11,127 (17,683 prioritized 1 kb regions) enhancer anchors in total; 7,341 (12,385 prioritized 1 kb regions) enhancer anchors are contacted by one promoter anchor with a maximum of 21 promoter anchors (15 using prioritized enhancer regions) sharing the same enhancer.

### E-P HiChIP prioritization

In order to reduce experimental artifacts in the context of our EPINs, we developed a specific prioritization method. This prioritization start by normalizing the data assuming, as most used capture-C normalizations (ICE ^52^, Vanilla, or KR ^53^) that all biases (e.g. GC content, number of restriction sites, mappability, or in the case of HiChIP, immunoprecipitation bias) can be corrected together. For this normalization step, we assume that there is a specific bias per any 1 kb genomic loci (L_x_ for loci x; see **Figures S1A and S1B**). This bias causes the difference between a theoretical expected number of interactions (E_XY_ between loci *X* and *Y*) and the observed number of interactions (O_XY_ between loci *X* and *Y*). In this representation we can define a system of 9 equations involving three 1 kb loci in the promoter (exactly from TSS -1 kb to TSS +2 kb) and fifteen 1 kb loci on the enhancer side. This system of equations is then solved using Sequential Quadratic Programming (SQP) ^54^. The procedure is repeated in an overlapping window manner along the 15 kb of the enhancer, always against the target 1 kb of the promoter and its two 1 kb neighboring loci. Before the normalization step, we observed a different interaction pattern for interactions below 10 kb (**Figure S1C**) due, in part, to the contiguity of restriction-enzyme fragments or chromatin persistence length. As these interactions may also be a source of bias in the construction of a PPI network, we removed them from our study. We applied the normalization to the remaining interactions and observed a better correlation between genomic distance and interaction count (**Figures S1D**).

In order to compare with standard normalization procedure we applied the ICE normalization^52^ to our dataset (using TADbit^55^ 1 kb resolution; filtering bins with less than 100 di-tags - 75% of the genome lost even using a threshold 10 times below the recommended^53^). Because of the sparsity of the genomic matrix the normalization did not fully converge (ICE was not able to completely balance the average di-tag counts per bin^52^). Next we applied the following normalization to our loops dataset, with few modification in order to rescue as much signal as possible: 1-in the promoter site, as our definition of promoter is exact (TSS to TSS +1 kb), we corrected using the average of the two bins spanning over this 1 kb region 2-on the enhancer site, as most of the 1 kb loci were excluded by the normalization filter we also averaged the ICE bias over the whole region. Even with these modifications, only half the original data was recovered. However the correlation between genomic distance and number of interactions was significantly improved with respect to raw data. Overall, the correlation value observed with ICE was similar to the one measured for our normalization (**Figure S1E**). We believe however that, for this dataset and for our methodology, our normalization procedure represents an improvement as it considers the exact promoter regions (not partially overlapping 1 kb bins) and minimizes the loss of promoter-enhancer data.

The normalized profile of interactions was finally used to prioritize most interacting 1 kb loci on the 15 kb enhancer (**Figure S1F**). The selected 1 kb regions are referred to as prioritized enhancer regions.

### DNA binding motifs

DNA binding motifs were retrieved from JASPAR (Fornes et al. 2019), an open-access database of curated, non-redundant binding profiles of DBPs (a.k.a. motifs) stored as position frequency matrices (PFMs). To detect the binding motifs, we used FIMO from the MEME-suite software (Grant et al. 2011), with p-value <= 1e-4 and q-value <= 5e-2 cutoffs. JASPAR contains **810 DNA binding motifs of 640 proteins** that overlap the E-P contacts identified with HiChIP.

### Gene expression data

We assayed RNA sequencing (RNA-seq) in the cell line LNCaP and LHSAR for two replicates using the VIPER pipeline as previously described ^11^, and fragments per kilobase of transcript per million mapped reads (FPKM) values were calculated for 20,114 RefSEQ genes. Genes with expression levels above the threshold of 0.003 in both replicates were considered in the entire analysis (**Figure S2**). Depending on the dataset, this expression lower-bound may be modified in different use cases, for instance based on specific insights or based on a differential analysis between conditions. In this work, we used FPKM instead of more direct measures as we set our threshold very low and did not want to enrich our dataset with very long, virtually unexpressed, transcripts.

### Protein-protein interaction network

We obtained protein-protein interactions (PPIs) from the Integrated Interactions Database (IID) ^56^. To better contextualize the interactome information, we combined the annotations of the PPIs from IID database with the LNCaP gene expression data. As for the IID annotations, we applied the following selection criteria. First we selected interactions annotated as “experimental” in the “*evidence type*” field and identified by at least two independent biological assays reported in the “*methods*” field. Then, we filtered only for interactions in the *prostate* or in *prostate cancer* cells and between *nuclear* proteins. Finally, we retain proteins whose gene expression levels were FPKM > 0.003 in both replicates (this cut-off removes ∼30% of the genes). In total, 14,221 proteins from a pool of 20,111 human protein coding genes meet the gene expression criteria. The combination of the above filtering criteria (gene expression, using only nuclear, prostate cancer or prostate and experimentally by 2 methods) resulted in an unweighted network of **31,944 prostate-specific nuclear PPIs among 4,295 proteins** ^56^.

Similarly, for the comparison with the LHSAR cell line we reconstructed the PPI interaction networks with PPIs from the same database (IID) having the following annotation criteria: “experimental” in the “*evidence type*” field and identified by at least two independent biological assays reported in the “*methods*” field. Then, we filtered only for interactions in the *prostate* cells and between *nuclear* proteins. Finally, we retain PPIs between proteins whose LHSAR gene expression levels were FPKM > 0.003 in both replicates. In total 29,316 PPIs representing 4,363 proteins were used for the EPIN reconstruction in the LHSAR cell line. Jaccard Index between the two resulting PPIs between LNCaP and LHSAR is 0.852.

### The PENGUIN pipeline

We set up graph-based approach, called Promoter-ENhancer-GUided Interaction Networks (PENGUIN), to reconstruct individual networks of protein interactions that might occur between one promoter (P) and its contacting enhancers (E), that we call E-P protein-protein Interaction Networks (EPINs). To reconstruct the EPINs, PENGUIN integrates information about chromatin contacts, protein-DNA binding, and protein-protein interactions (PPIs). For the case under study in this work (prostate cancer, PrCa), chromatin contacts information comes from H3K27Ac HiChIP of LNCaP cells (4,314 promoters and 5,789 enhancer regions; see **Methods**, “Conformation capture and E-P interactions’’), protein-DNA binding information ^53, 54^ comes from the JASPAR database (810 DNA binding motifs of 640 proteins; see Methods, “DNA binding motifs”), and PPIs information comes from the IID database (31,944 prostate-specific nuclear PPIs among 4,295 proteins; see Methods, “Protein-protein interaction network”) further filtered using LNCaP RNA-seq data (see Methods, “Gene expression data”).

The reconstruction of EPINs follows these steps: for each E-P contact, (1) DNA binding motifs are detected in the corresponding sequences of promoter and enhancer regions; (2) a subnetwork of PPIs is selected containing all promoter-bound proteins, all enhancer-bound proteins, and all their intermediate interactors, with a maximum of 1 intermediate node between enhancer and promoter bound DNA binding proteins; (3) intermediate interactors are discarded if they only connect promoter-bound proteins or enhancer-bound proteins. Using the provided PrCa information, PENGUIN reconstructed 4,314 EPINs consisting of a total of 9,141 PPIs among 885 proteins of which 751 are intermediate proteins linking promoter-bound and enhancer-bound proteins.

### Node centrality measures

In several analyses we employed two measures of node centrality, namely betweenness and degree. **Betweenness** is a measure of centrality in a graph based on shortest paths. For every pair of nodes in a connected graph, there exists at least one shortest path between the vertices such that either the number of edges that the path passes through is minimized. The **degree** of a node in a network is the number of connections it has to other nodes; the degree distribution is the probability distribution of these degrees over the whole network.

### Clustering EPINs

We defined EPIN clusters by taking into account their edge content. Each edge consists of an individual pairwise PPI as defined previously. We collected the full universe of edges using all existent edges between all promoter EPINs (the union graph). Then we computed the distance between EPINs by counting the number edges shared over the total number of edges in our predefined universe of edges. Finally we performed clustering using this distance matrix from all possible combinations of EPIN pairs. The clustering was performed using Ward’s linkage method. Each leaf in the obtained cluster represents a promoter EPIN.

### Identifying enriched functional annotations in EPIN clusters

We performed two-sided Fisher’s exact tests on every single branch of the dendrogram representing the obtained hierarchical clustering. We evaluated the enrichment of any feature (CTCF binding sites by ChIP-seq, PrCa SNPs from curated GWAS, PrCa oncogenes) in the leaves under a branch of interest compared to those in the rest of the tree. For the enrichment in CTCF binding, we used CTCF peaks from an external dataset but in the same cell line (see **CTCF ChiP-Seq peaks).** We considered an EPIN to be CTCF-positive (CTCF+), if a CTCF peak was found in a 10 kb region around its promoter and around 10 kb of at least one of its enhancer regions.

For the GWAS feature, we require the presence/overlap of a PrCa-associated SNP (see **Genome-wide association data**) in at least one of the enhancers of an EPIN. Two-sided Fisher’s exact tests were used to calculate the odds ratio (OR) and enrichment p-values for presence of PrCa annotations within the identified clusters.

### Druggability information

We extracted information for target druggability from DrugBank ^57^. The use of each drug was obtained from the Therapeutic Target Database ^58^. We annotated each protein node that is a target of drugs that are assigned as Approved or under Clinical Trials (Phase 1, 2, 3) or Investigable for Prostate Cancer, as PrCa druggable.

### CTCF ChiP-Seq peaks

CTCF ChIP-seq peaks for LNCaP cell line were retrieved from ENCODE51 project (https://www.encodeproject.org/) for the same Genome assembly, hg19 (GEO references: *GSM2827202* and *GSM2827203*). Overlaps of the CTCF binding sites with enhancer and promoter anchors allowed a 10 kb gap between them. Since CTCF ChiP-seq peaks for LHSAR cell line were not available in ENCODE, we retrieved from ChIP Atlas (https://chip-atlas.org/) two distinct sets (GEO references: *GSM2825573* and *GSM2825574*) of CTCF peaks (of same Genome assembly hg19) for prostate epithelial cells at a q-value of 1e-10 (Table 3). We used these two sets independently and in concatenation when comparing the clustering results between LNCaP and LHSAR. These narrow peaks were mapped on the enhancer regions using the python package *PyRanges* (see “E-P contacts” section). For both cases, LNCaP and LHSAR, the narrow peaks were considered as the CTCF binding sites.

### PrCa SNPs

To explore enrichment of SNPs associated to PrCa across the identified clusters, and to identify the SNP paths, we used the previously reported 95% credible set^11^ from fine-mapping 137 previously-associated PrCa regions using a Bayesian statistical method PAINTOR ^59^ employing the largest PrCa genome-wide association studies (GWAS) (N = 79,148 cases and 61,106 controls)^60^. This set was composed of 5,412 distinct SNPs (rsid). We will refer to these as PrCa SNPs. Note that this set also includes SNPs that do not reach genome-wide-filters of p-value significance. We illustrate the location of the associated PrCa regions and number of PrCa SNPs in **Figure S11**. We did not find a significant correlation between the number of PrCa SNPs in the regions and the number of PrCa SNPs we prioritized in this work (Pearson r=0.2, p-value=0.06 and Pearson r=0.1, p-value=0.3 for **Tables S10** and **S11**, respectively). We mapped the SNP location to prioritized enhancer regions anchor locations with a window of 10 kb. 518 out of 5,412 overlap our prioritized enhancer regions; 18 of them overlap our promoter regions. In total 218 prioritized enhancers and 14 promoters overlap a PrCa SNP.

### SNP paths (PrCa SNPs in enhancer binding motifs)

A path in a network is a sequence of edges joining a sequence of nodes. We detected PrCa SNPs located in the DNA binding motifs in the enhancers, and identified the corresponding SNP paths (linked edges and nodes) for each EPIN promoter. For SNP paths analyses and the web-browser, we used all PrCa SNPs in the 95% credible set. There were 36 PrCa SNPs falling in enhancer binding motifs across clusters 3, 4, 5, 6, 7, 8. To report the most interesting cases in the Tables and Results, we used the subset of those passing genome-wide significance of p-value for PrCa association < 5e-8. There were 15 PrCa SNPs falling in enhancer binding motifs across clusters 3, 5, 6, 7, 8.

### SNP paths (PrCa SNPs in intermediate proteins)

We detected PrCa SNPs falling within genes that encode for intermediate nodes, and identified the corresponding SNP paths (linked edges and nodes) for each EPIN promoter. For SNP paths analyses and the web-browser, we used all PrCa SNPs in the 95% credible set..

### PrCa enrichment using GWAS Catalog and comparison with other diseases

This analysis had two aims: 1) explore whether we could replicate our finding and identify the GWAS enriched cluster using a different source for the PrCa associated SNPs using SNPs extracted from the GWAS catalog instead of fine-mapped SNPs; 2) to compare the GWAS signal for different diseases. We estimated enrichment of SNPs overlapping the enhancers in each of the identified clusters by exploring the NHGRI GWAS Catalog associations ^61^. First, we retrieved GWAS data and filtered the traits according to their “umlsSemanticTypeName” as defined in DisGeNet database ^62^ to one of the following: "Mental or Behavioral Dysfunction", "Neoplastic Process", "Disease or Syndrome", "Congenital Abnormality; Disease or Syndrome", "Disease or Syndrome; Congenital Abnormality", "Disease or Syndrome; Anatomical Abnormality". We considered only traits with at least 10 genome-wide-significant SNPs (unadjusted p-value < 5e-8). We mapped the SNP location to prioritized enhancer anchor locations with a window of 10 kb. 104 diseases had SNPs overlaps and 17 of them have more than 10 SNP overlapping (**Table S5).** For each cluster, we tested enrichment of disease-associated SNPs using Fisher tests and considered significant p-value < 0.01 and OR > 1.

### Trans-eQTL hotspots

We retrieved trans-eQTLs reported in the largest meta-analysis with up to 31,684 blood samples from 37 eQTLGen Consortium cohorts in whole blood in ^22^. We grouped enhancers by collapsing when they were separated by less than 20 kb, thereby creating ‘enhancer clusters’. To qualify as a trans-eQTL hotspot, the enhancer clusters had to contain a SNP associated with at least 3 different genes. We quantified the normalized mutual information (NMI) between the hotspot-related enhancer clusters and our 8 EPIN clusters. In order to infer deviation from expected by chance and estimate an empirical p-value, we randomized 10 thousand times the association between each enhancer and its corresponding EPIN cluster and computed the NMI between each randomized EPIN clustering and the observed hotspot-related enhancer clustering. Additionally, we checked if a given cluster was significantly enriched in trans-eQTL hotspots. For this purpose we applied a Fisher test to our pool of enhancers comparing the two contingencies, inside/outside a given cluster, and inside/outside a trans-eQTL hotspot.

### Super-enhancer-like regions

We defined enhancer hotspots as groups of enhancers separated by less than 15 kb, and identified 3,752 enhancer hotspots using *bedtools cluster*.

### Oncogenes Gene list

We used a previously identified list of 122 Genes ("PrCa_GeneList_Used.csv") known to be somatically mutated in PrCa oncogenesis (37 out of 4,314 promoters considered). As previously described ^11^, the 122 oncogenes are a set of prostate cancer–genes curated from three large-scale PrCa studies that show evidence of somatically acquired mutations, at both localized and advanced prostate cancer, known and recurrently altered in localized prostate cancer and metastatic prostate cancer.

### Enriched edges within each cluster

Two-sided Fisher’s exact tests were used to compute odds ratios and p-values of the edges and nodes in the eight different clusters. Specifically, each edge or node was tested for presence/absence in a cluster compared to all others. Therefore, one edge or node can be enriched in one or more than one cluster, it cannot be enriched in all clusters.

### Significantly central intermediate nodes within each cluster

We computed protein importance for each cluster in terms of two network centrality measures: betweenness and degree. For each protein we obtain both betweenness and degree specificity ratios in order to equitably quantify internal protein centrality differences between the clusters. For each of the found clusters we independently estimated the specificity of the observed protein centrality measures (“Betweenness” and “Degree”). For a given protein (P_i_) in a particular cluster (C_j_), we define the specificity as the ratio between the mean centrality value of P_i_ inside the fraction of networks belonging to C_j_ ; divided by the mean centrality value of P_i_ for the fraction of networks outside of the cluster C_j_.

*Specificity ratio* (P_i_, C_j_) = (mean (Pi centrality in C_j_ networks) + 1) / (mean (P_i_ centrality in non C_j_ networks) + 1)

We assessed protein specificity ratio significance for each cluster upon random network cluster generation. Aiming to assess the significance of the different specificity ratios for the proteins within each cluster, we developed a significance analysis test based on random cluster subsamplings. In order to compute the significance of a given protein specificity ratio (P_i_) within a particular cluster of analysis (C_j_), we performed 1000 random network samplings to produce random network clusters containing the same amount of networks as the real cluster being analyzed (i.e. if the real cluster contains 100 networks, the random clusters generated will contain 100 random networks out of the 4,314 clustered networks). Within each of those 1000 random clusters, we compute the corresponding protein specificity ratios, with the p-value representing the probability of finding the protein specificity ratio to be higher or equal to the real value computed for the particular cluster of interest (C_j_).

We also performed Fisher tests to assess enrichment for the presence of the node in the cluster (Fisher test p-value < 0.01). EP300 was excluded from the enrichment test as the presence of that node was not significantly enriched (Fisher test p-value < 0.01). 22 proteins (SMAD2, KAT5, NCOR2, MAPK8, SMAD4, CREBBP, CTNNB1, PGR, HDAC3, HDAC2, GSK3B, UBA52, UBE2I, JUND, PIAS1, XRCC5, CDK6, XRCC6, MAPK1, FOS, HIF1A and MAPK3) were found to be significantly specific for both betweenness and degree ratios (p-value < 0.01 for both centrality measures and over-represented in this cluster using Fisher tests) and used as input for the functional gene set enrichment analysis presented as **Table S9**).

We provide the full results of the centrality significance analysis for each cluster in github:

https://github.com/bsc-life/penguin_software/tree/main/Protein_Significance_analysis

### Functional gene set enrichment analysis

Functional enrichment analysis was performed using the g:GOST module from g:Profiler, a web tool to perform simultaneous gene set enrichment analysis across multiple biomedical databases ^23^. We query the web service using the R implementation available from gprofiler2 package. g:GOST performs cumulative hypergeometric tests of an input geneset against preprocessed database-specific gene sets.

The code for this analysis is available as a Jupyter Notebook that can be accessed in github:

https://github.com/bsc-life/penguin_software/tree/main/gProfiler_GSEA/Supplementary_Tables_5_7_9_and_Significantly_Central_Protein_Enrichment_Analysis.ipynb

We set alternative backgrounds for the gene set enrichment analysis, depending on the analysis. For the analysis presented as **Table S5**, where we run the web service to test functional enrichment of the genes associated to the promoter networks from cluster 8, the background is set to the 4,314 genes associated with the clustered EPINs. For the analysis presented as **Table S7**, where we test for general functional enrichment of all different proteins forming the EPINs, we run the web service considering only annotated genes for the statistical domain scope. Finally, for the analysis presented as **Table S9**, where we test the functional enrichment of the significantly central (p-value < 0.01 for both degree and betweenness centrality) proteins of networks from GWAS+ cluster, the background is formed by the very limited set of 751 unique intermediate proteins forming the EPINs. We additionally provide, within the very same Jupyter Notebook, comparative dot plots presenting the functional enrichment analysis of significantly central proteins of each cluster under **Table S9** setting.

Reported adjusted p-values correspond to Benjamini-Hochberg correction for multiple testing, with adjusted p-values ≤ 0.05 considered to be significant. Gene set enrichment analysis results are provided for KEGG pathways, Reactome, Gene Ontology, Wikipathways, TRANSFAC, miRTarBase, Human Protein Atlas, CORUM and Human Phenotype Ontology. For the enrichment analysis of significantly specific proteins of the GWAS+ cluster, we provided as input the 22 previously described proteins. For the enrichment analysis of the GWAS+ cluster, we provided as input all genes associated with the EPIN promoters in cluster GWAS+.

### Differential Gene Expression

We integrated data from EPIN promoters with differential gene expression (DE) from two sources. DE analysis on prostate cancer tumor versus normal was downloaded from GEPIA: http://gepia2.cancer-pku.cn/#degenes, which use the TCGA and GTEx projects databases to compare gene expression between tumor and normal tissues under Limma, both under and over expressed. We used the default thresholds of log2FC of 1 and qvalue cut-off of 0.01. These data covered 84 out of 885 genes encoding for intermediates in PENGUIN and 413 out of 4,314 promoter EPINs. DE analysis of RNA-Seq on LHSAR (an immortalized prostate epithelial line overexpressing androgen receptor) versus LNCaP was performed as previously described. Briefly, RNA-seq data were processed using the VIPER pipeline ^63^. Reads were aligned to the hg19 human genome built with STAR. FPKM values were calculated with Cufflinks for 20,114 RefSEQ genes included in the VIPER repository. Differential expression analysis was performed with the DESeq2 R package ^64^. 15,650 genes with DE data covered 884 of the 885 genes encoding for intermediates in PENGUIN and 3,286 genes out of 4,314 promoter EPINs.

We annotated whether the EPIN promoters themselves and the genes encoding the intermediate proteins in our data were DE using either of the two databases. We considered as DE those genes passing |log2 fold change| > 1 and adjusted p-value <= 0.01. For the LNCAP/LHSAR dataset, we could compute a Fisher test of enrichment of differentially expressed genes encoding for intermediate proteins within each EPIN promoter versus within the SNP paths (we could not compute this for the GEPIA since we did not have the full dataset of covered genes). The genes that were not passing these filters were considered non-DE and the genes not covered by the two datasets were excluded from the enrichment analysis described next. For each EPIN we calculated the fraction of DE intermediates within the SNP paths and we estimated the enrichment of those compared to the fraction of DE intermediates in the full EPIN network.

To find the enrichment of DE genes in SNP paths (PrCa SNPs in intermediate proteins) compared to those in the entire EPIN, we computed as enrichment the ratio of Fraction1 / Fraction2, where:

Fraction1 = (number of DE intermediates within SNP paths) / (number of covered intermediates within SNP paths), and

Fraction2 = (number of DE intermediates the EPIN) / (number of covered intermediates in the EPIN).

We report the EPIN genes passing enrichment (“**enrichment_DE_deseq_SNP.bs.DBP.path**”) > 1.

### pQTL look-up

We downloaded summary statistics with genome-wide association between SNPs and 4907 proteins reported in the deCODE study ^65^ and annotated with pQTL association the SNPs we identified falling either in enhancer binding sites or in node genomic locations. The deCODE pQTL summary statistics data contained information on 4,907 proteins and 186 (201 PrCa SNPs out of the 213 PrCa SNPs we looked up were in the data and 186 also matched by alleles). 808 out of the 4,314 genes promoters ("EPIN_promoters") and 278 out of the 885 gene intermediates (in total 997 out of 4,918 genes promoters and coding for intermediates in our networks) have information on associations with their respective coded proteins covered by the pQTL deCODE data.

### Gene dependency and gene effect metrics

*Gene Effect* and *Gene Dependency* metrics were downloaded from the DepMap portal (https://depmap.org/portal/). We used both the RNAi ^66^ and CRISPR ^67^ datasets.

### Data Availability

RefSeq hg19 from UCSC Genome Browser is available at the following URL: http://genome.ucsc.edu/cgi-bin/hgTables?hgsid=694977049_xUU5i1QkIJ50dj5miBt9wkAYuxN3&clade=mammal&org=&db=hg19&hgta_group=genes&hgta_track=knownGene&hgta_table=knownGene&hgta_regionType=genome&position=&hgta_outputType=selectedFields&hgta_outFileName=knownGene.gtf

All EPINs and related statistics can be downloaded through the PENGUIN web service at https://penguin.life.bsc.es/

All the raw listed in **Table 3**, as well as the corresponding processed and metadata for LHSAR and LNCaP related to H3K27ac (HiChIP) and RNAseq have been deposited in GEO. CTCF ChIP-Seq data used in this work comes from ENCODE^51^ with references GSM2827202, GSM2827203 for LNCaP and GSM2825573, GSM2825574 for the human epithelial cells or prostate that we use to infer CTCF-bindings in LHSAR GSM2825573, GSM2825574

### Code Availability

Source code of the related to the PENGUIN protocol is available at github: https://github.com/bsc-life/penguin_software

Source code of the related to the PENGUIN web service is available at github: https://github.com/bsc-life/penguin_analytics

R (v.4.2.0) and Python were extensively used to analyze data and create plots. biomart / ensembl from biomaRt package Ensembl hg19 data for overlaps of SNPs with intermediates.

### Competing interests

None of the authors have competing financial or non-financial interests.

## Notes

https://penguin.life.bsc.es/

### Competing Interest Statement

The authors have declared no competing interest.

### Summary of Updates

We extensively revised the abstract, introduction, and conclusions sections to address this concern (Figure 1). To ensure a comprehensive analysis, we utilized a benign human prostate epithelial cell line (LHSAR) as a baseline for comparison with LNCaP cells. We performed HiChIP experimental data generation specifically for LHSAR cells and applied the PPI clustering procedure to explore functional relationships within the acquired data. Our analysis did not reveal cluster enrichment in GWAS and CTCF within the benign LHSAR prostate control cell line (Figure 2). The experimental data have been deposited in GEO and in https://github.com/bsc-life/penguin_software. We provided further evidence on the clustering procedure and properties (Figure 2). We performed additional statistics on the PPIs of different cells to support our claim that information in protein-protein networks allows us to identify disease-specific networks. We quantified the overlap between PPI networks and demonstrated that the interactions are highly cell-specific (Figure S6). We repeated our clustering analysis, based solely on the list of enhancer IDs in each EPIN. Our results showed that the use of intermediate PPI networks increases the number of identified oncogenes, indicating the relevance of the PPI network for EPIN classification and its association with phenotype (Figure S7). We validated the top candidates reported in Table S10 and Table S11 through previously pooled genome-wide CRISPR/Cas9 knockout and RNAi screens conducted in prostate cancer LNCaP cells (Figure 4). In addition to MYC, we included more examples, such as GATA2 and CASC11, to further support our findings (Figure S8).

