## Supplementary Materials for "The PENGUIN approach to reconstruct protein interactions at enhancer-promoter regions and its application to prostate cancer"

1 Istituto Italiano di Tecnologia, Via Morego, 30 16163 Genova, Italy.
2 Barcelona Supercomputing Center, Plaça Eusebi Güell, 1-3, 08034 Barcelona, Spain
3 Present: Josep Carreras Leukaemia Research Institute, Badalona, Barcelona, Spain
4 Department of Medical Oncology, The Center for Functional Cancer Epigenetics, Dana Farber Cancer Institute, Boston, MA 02215, USA.
5 ICREA - Institució Catalana de Recerca I Estudis Avançats, Pg. Lluís Companys 23, 08010 Barcelona, Spain
6 Sapienza University Rome, Biology and Biotechnologies Department C. Darwin, ologV.le Aldo Moro 5, 00185

** these authors contributed equally to this work*
*[#] to whom correspondence should be addressed*

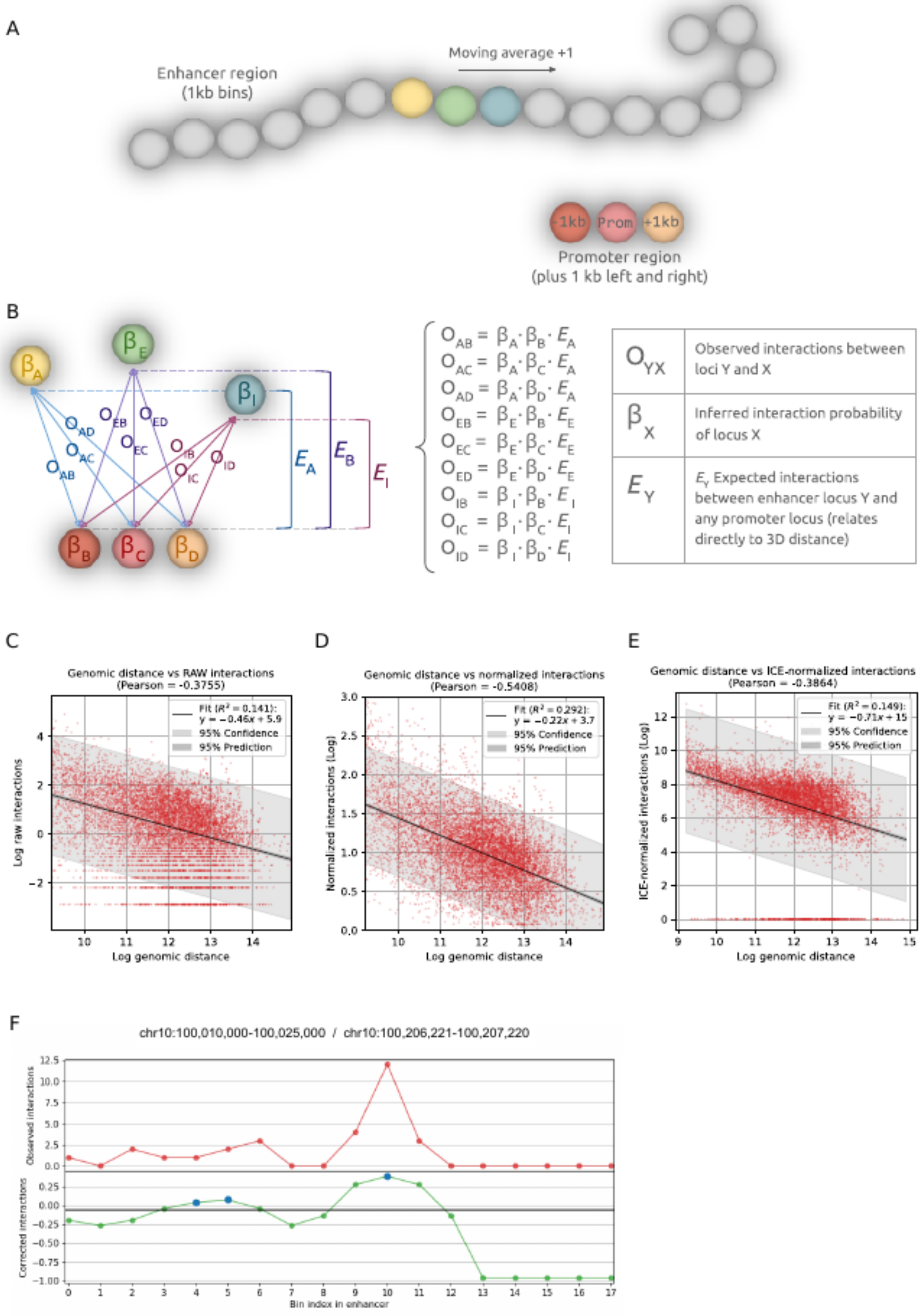
A
Moving average +1
Enhancer region
(1kb bins)
-1kb
Prom
+1kb
Promoter region
(plus 1 kb left and right)
B
O_AB = β_A · β_B · E_A
O_AC = β_A · β_C · E_A
O_AD = β_A · β_D · E_A
O_EB = β_E · β_B · E_E
O_EC = β_E · β_C · E_E
O_ED = β_E · β_D · E_E
O_IB = β_I · β_B · E_I
O_IC = β_I · β_C · E_I
O_ID = β_I · β_D · E_I
O_YX Observed interactions between loci Y and X
β_X Inferred interaction probability of locus X
E_Y E_Y Expected interactions between enhancer locus Y and any promoter locus (relates directly to 3D distance)
C
Genomic distance vs RAW interactions
(Pearson = -0.3755)
Fit (R² = 0.141): y = −0.46x + 5.9
95% Confidence
95% Prediction
Log raw interactions
Log genomic distance
D
Genomic distance vs normalized interactions
(Pearson = -0.5408)
Fit (R² = 0.292): y = −0.22x + 3.7
95% Confidence
95% Prediction
Normalized interactions (Log)
Log genomic distance
E
Genomic distance vs ICE-normalized interactions
(Pearson = -0.3864)
Fit (R² = 0.149): y = −0.71x + 15
95% Confidence
95% Prediction
ICE-normalized interactions (Log)
Log genomic distance
F
chr10:100,010,000-100,025,000 / chr10:100,206,221-100,207,220
Observed interactions
Corrected interactions
Bin index in enhancer

**Figure S1: HiChIP promoter-enhancer loop prioritization. (A)** Schematic representation of a promoter enhancer loop, where the promoter is represented by a 1kb bead surrounded by two neighbor beads, and the enhancer by fifteen 1kb beads. **(B)** Representation of the parameters taken into account to compute the expected number of interactions between a 1 kb loci from the enhancer and the 1kb loci from the promoter. **(C)** Correlation between the genomic distance between enhancer and promoter and the number of interactions. **(D)** Same as (C) but with data normalized by the strategy explained in B, (**E**) Same as (C) but with data normalized by ICE **(F)** Example of profile of raw interactions along an enhancer (top), and normalized interactions (bottom). Only beads highlighted in blue in the bottom plot would be used (*prioritized*) in the PENGUIN analysis.

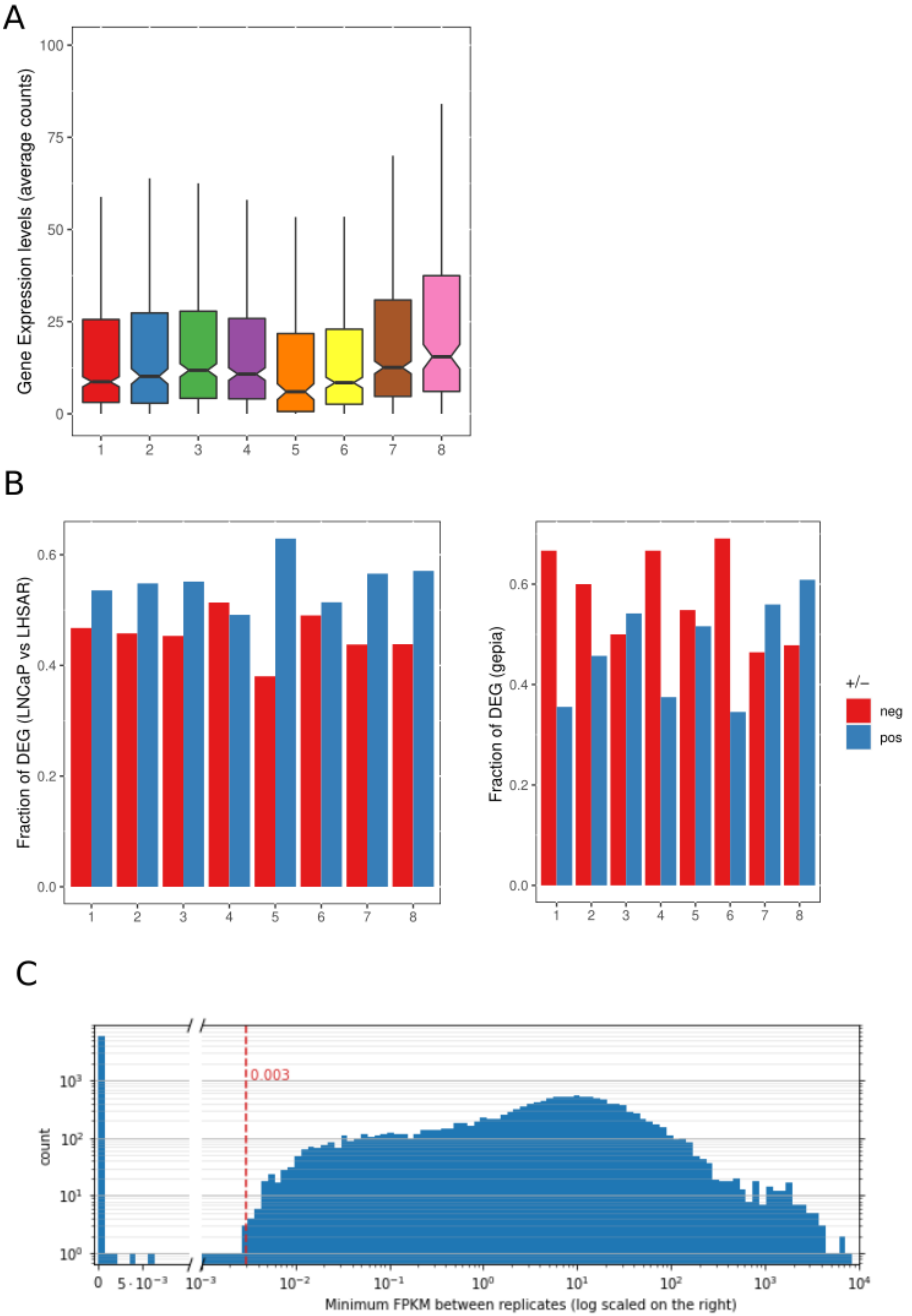
A
Gene Expression levels (average counts)
100
75
50
25
0
1 2 3 4 5 6 7 8
B
Fraction of DEG (LNCaP vs LHSAR)
0.6
0.4
0.2
0.0
1 2 3 4 5 6 7 8
Fraction of DEG (gepia)
0.6
0.4
0.2
0.0
1 2 3 4 5 6 7 8
+/−
neg
pos
C
0.003
count
$10^3$
$10^2$
$10^1$
$10^0$
0
$5 \cdot 10^{-3}$
$10^{-3}$ $10^{-2}$ $10^{-1}$ $10^0$ $10^1$ $10^2$ $10^3$ $10^4$
Minimum FPKM between replicates (log scaled on the right)

**Figure S2: General statistics on gene expression analysis. (A)** Gene expression per cluster and per EPIN's representative gene. **(B)** Fraction of differentially expressed promoter EPINs (DEG) in each cluster. Positive DEG in blue, negative DEG in red. **(C)** Distribution of expression, showing, per gene, the minimum value between the two RNA-seq replicates (methods). The red dotted line represents the threshold to consider a gene to be expressed.

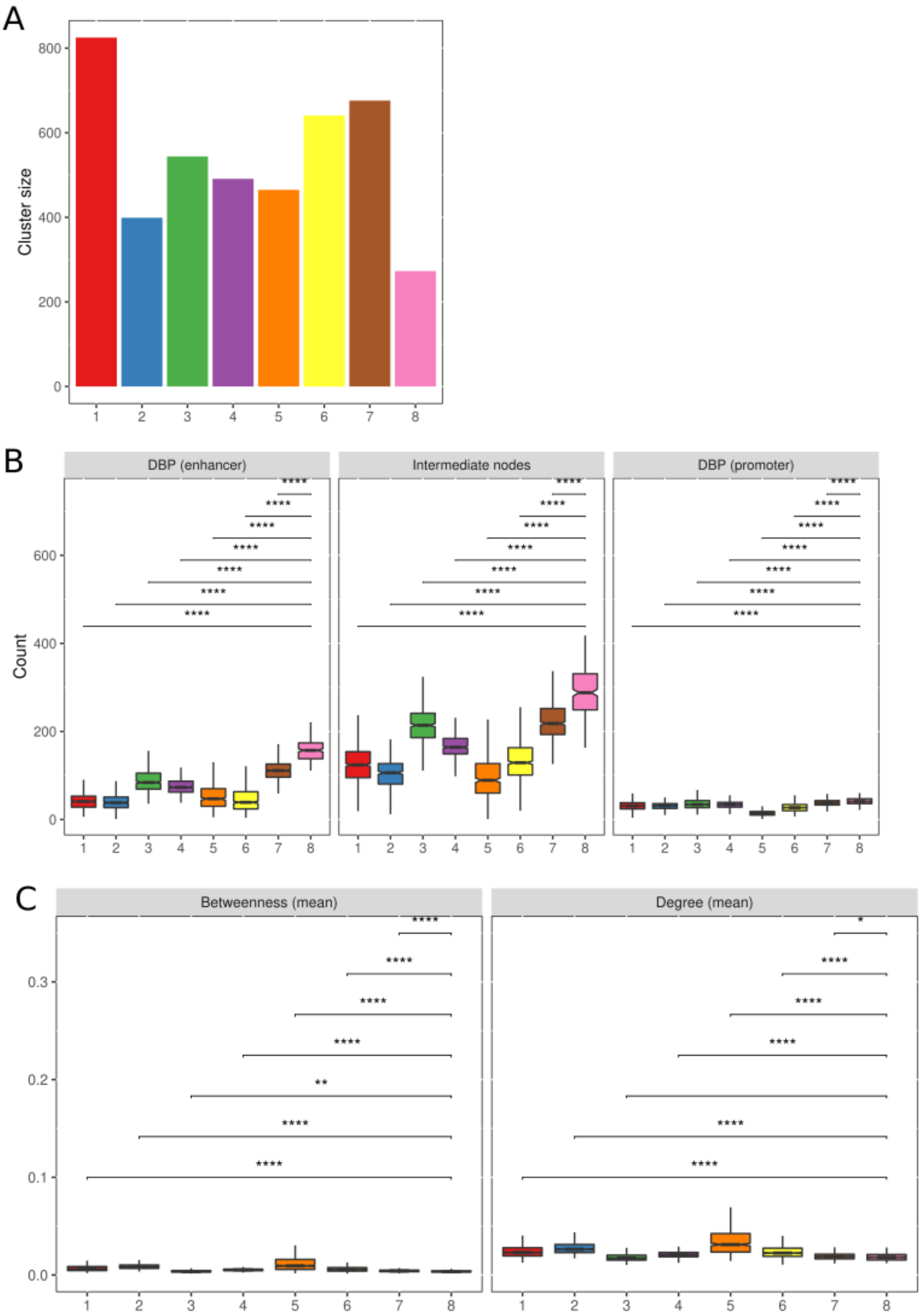
A
Cluster size
0
200
400
600
800
1 2 3 4 5 6 7 8
B
DBP (enhancer)
Intermediate nodes
DBP (promoter)
Count
0
200
400
600
C
Betweenness (mean)
Degree (mean)
0.0
0.1
0.2
0.3

**Figure S3: Descriptive statistics on EPIN clusters. (A)** number of EPINs per cluster, each EPIN is composed of one promoter and at least one enhancer. **(B)** Total number of DNA binding proteins (DBPs) potentially bound to enhancers (left), intermediate nodes from the PPI network between the promoter and its enhancers (center), and DBPs potentially bound to promoter (right), per EPIN cluster. **(C)** Centrality measures on intermediate nodes of EPINs per cluster. Namely betweenness (left) and degree (right); stars on the top indicate the degree of significance after a t-test test (*: p-value<0.05; **: p-value<0.01; ***: p-value<0.001; ***: p-value<0.0001).

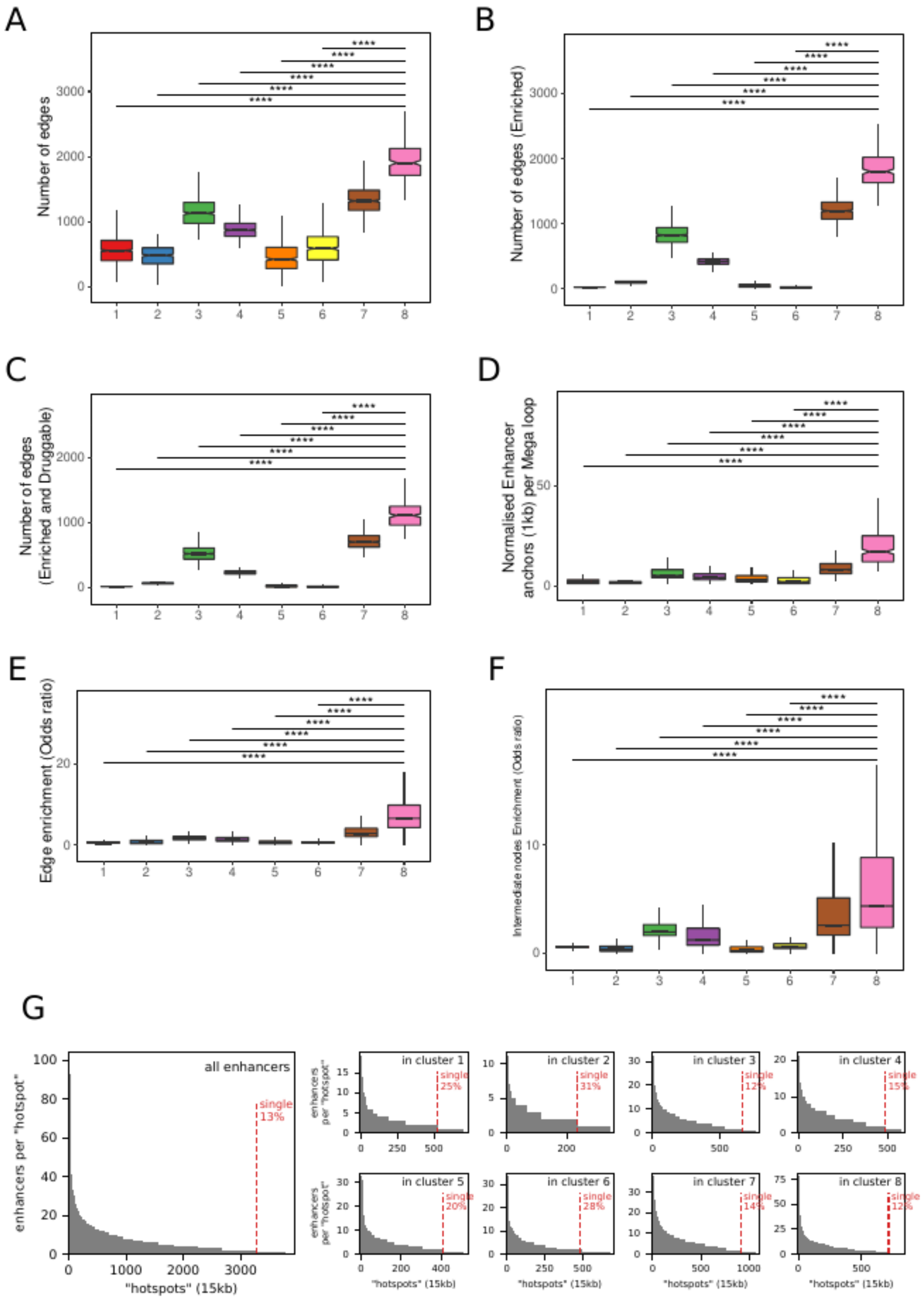
A
Number of edges
B
Number of edges (Enriched)
C
Number of edges
(Enriched and Druggable)
D
Normalised Enhancer
anchors (1kb) per Mega loop
E
Edge enrichment (Odds ratio)
F
Intermediate nodes Enrichment (Odds ratio)
G
enhancers per "hotspot"
"hotspots" (15kb)
all enhancers
single
13%
in cluster 1
single 25%
in cluster 2
single 31%
in cluster 3
single 12%
in cluster 4
single 15%
in cluster 5
single 20%
in cluster 6
single 28%
in cluster 7
single 14%
in cluster 8
single 12%
enhancers per "hotspot"

**Figure S4: Statistics on EPIN edges. (A)** Number of edges per EPINs grouped by clusters (boxplots). **(B)** Same with enriched edges (Fisher enrichment test in one cluster with respect to the others, see **methods**). (C) Same as B filtered by edges containing a druggable (see methods) target protein. (D) . (E). (F). Significance levels depicted represent the same as in **Figure S3**. (G) Number of prioritized enhancers per enhancer hotspots. Hotspots are defined as groups of enhancers separated by less than 15kb. Dotted red line shows the proportion of enhancers that are isolated. The different panels show enhancers in the whole genome (left), and in each of our 8 defined clusters (smaller panes on the right).

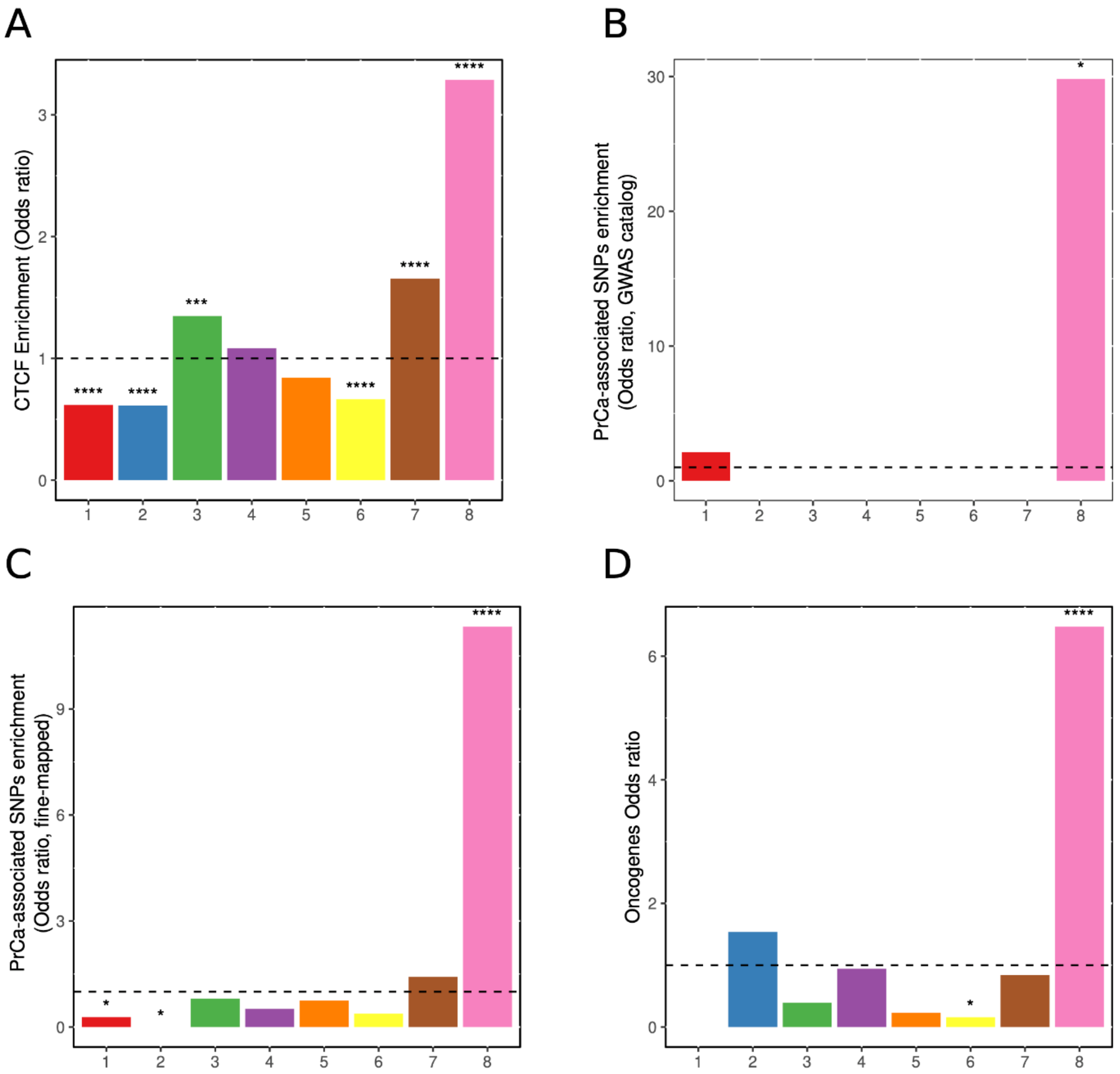


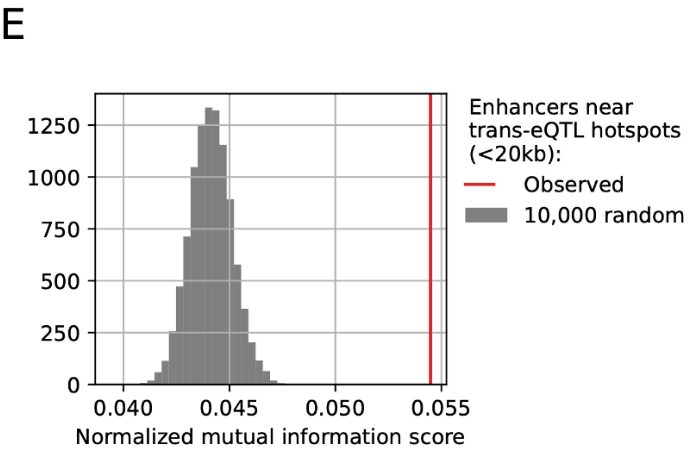


F

| Cluster number | Odd-ratio | p-value |
|---|---|---|
| cluster 1 | 1.1 | 0.65 |
| cluster 2 | 1 | 1 |
| cluster 3 | 0.8 | 0.35 |
| cluster 4 | 1.2 | 0.38 |
| cluster 5 | 1.2 | 0.61 |
| cluster 6 | 0.7 | 0.2 |
| cluster 7 | 1.1 | 0.58 |
| cluster 8 | 0.8 | 0.42 |

**Figure S5: Enrichment of EPIN clusters in biologically relevant features. (A)** CTCF. **(B)** GWAS SNPs. **(C)** GWAS SNPs paintor. **(D)** Oncogene Odd ratio. In all panels, stars represent significance of a fisher test against all clusters (*: p-value<0.05; **: p-value<0.01; ***: p-value<0.001; ***: p-value<0.0001). (**E**) Relationship between trans-eQTL hotspots and the 8 clusters using the concept of normalized mutual information. We focused on enhancers derived from our EPINs, which were associated with trans-eQTL hotspots located within a proximity of less than 20kb. A relatively weak correlation coefficient of 0.0546 if found between the 8 clusters and the hotspots defined by their proximity to trans-eQTL hotspots. Randoms were generated by shuffling the association between enhancers and EPIN clusters. (**F**) We investigated whether a specific cluster exhibits a significant enrichment of trans-eQTL hotspots. For this employed a Fisher test, comparing two contingencies within our list of enhancers: those within or outside a given cluster, and those within or outside a trans-eQTL hotspot.

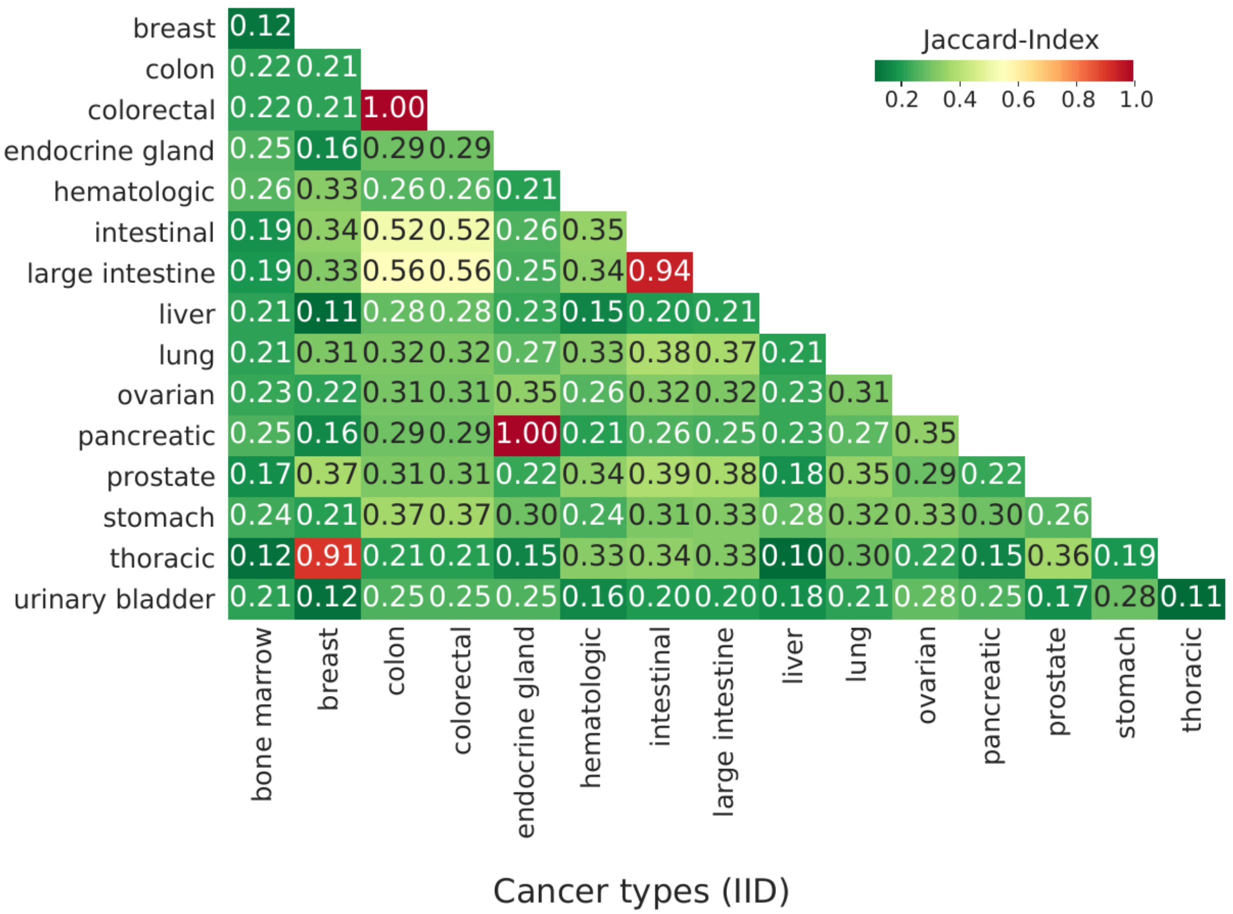


**Figure S6. PPI networks comparison.** Statistical analyses on PPIs across cancer cell types available at http://iid.ophid.utoronto.ca/. Using the Jaccard index we studied the overlap between PPI networks observing significant variations that were highly specific to each cell type. The results show that the PPIs used in PENGUIN vary significantly depending on the cell types of interest.

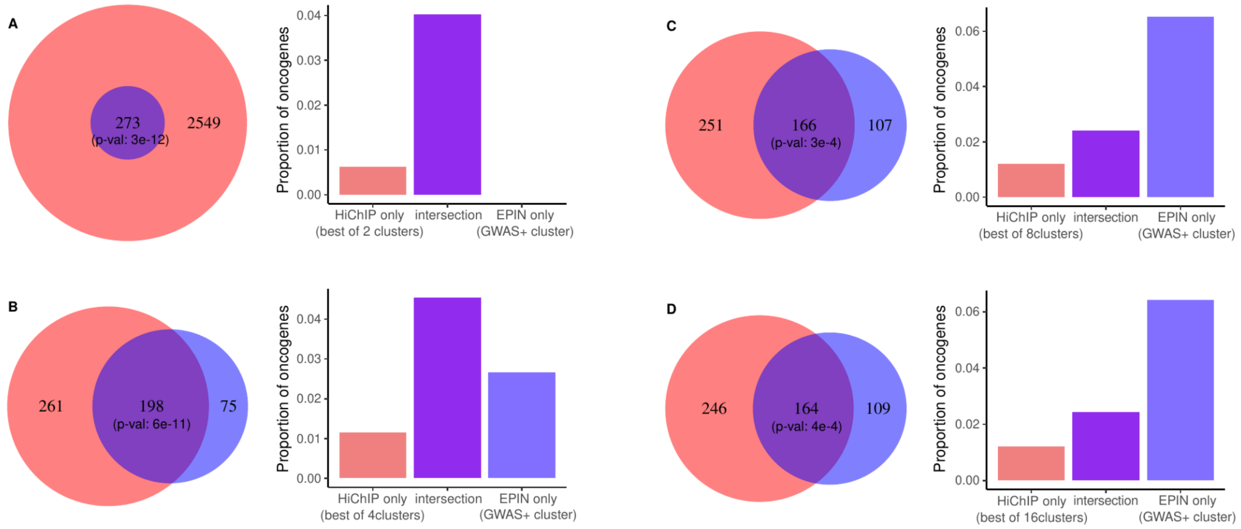


**Figure S7.** Comparison between clustering based on full EPINs (blue) and using only HiChIP data (no intermediate PPI network) (red). For each clustering strategy, only the cluster most enriched in PrCa SNPs and CTCF peaks is used in the comparison. The comparison is conducted in terms of the proportion of known PrCa oncogenes in the two sets, considering various cluster numbers within the red set (2, 4, 8, and 16 clusters), and only one cluster set (8 clusters for the blue set). Each panel (**A-D**) illustrates a Venn diagram showing the intersection (purple) between the red set and the blue set, and the corresponding fraction of oncogenes as a bar plot. The fraction of oncogenes that are unique to the red set ("HiChIP only") is consistently lower than the fraction of oncogenes that are unique to the blue set ("EPIN only"). Moreover, when compared with 8 and 16 clusters of the red set, the fraction of oncogenes of the "EPIN only" subset is higher than the intersection, indicating a relative gain in oncogenes retrieval when PENGUIN is employed. The significance of the intersection was estimated with a hypergeometric test considering the union of the two sets as the background.

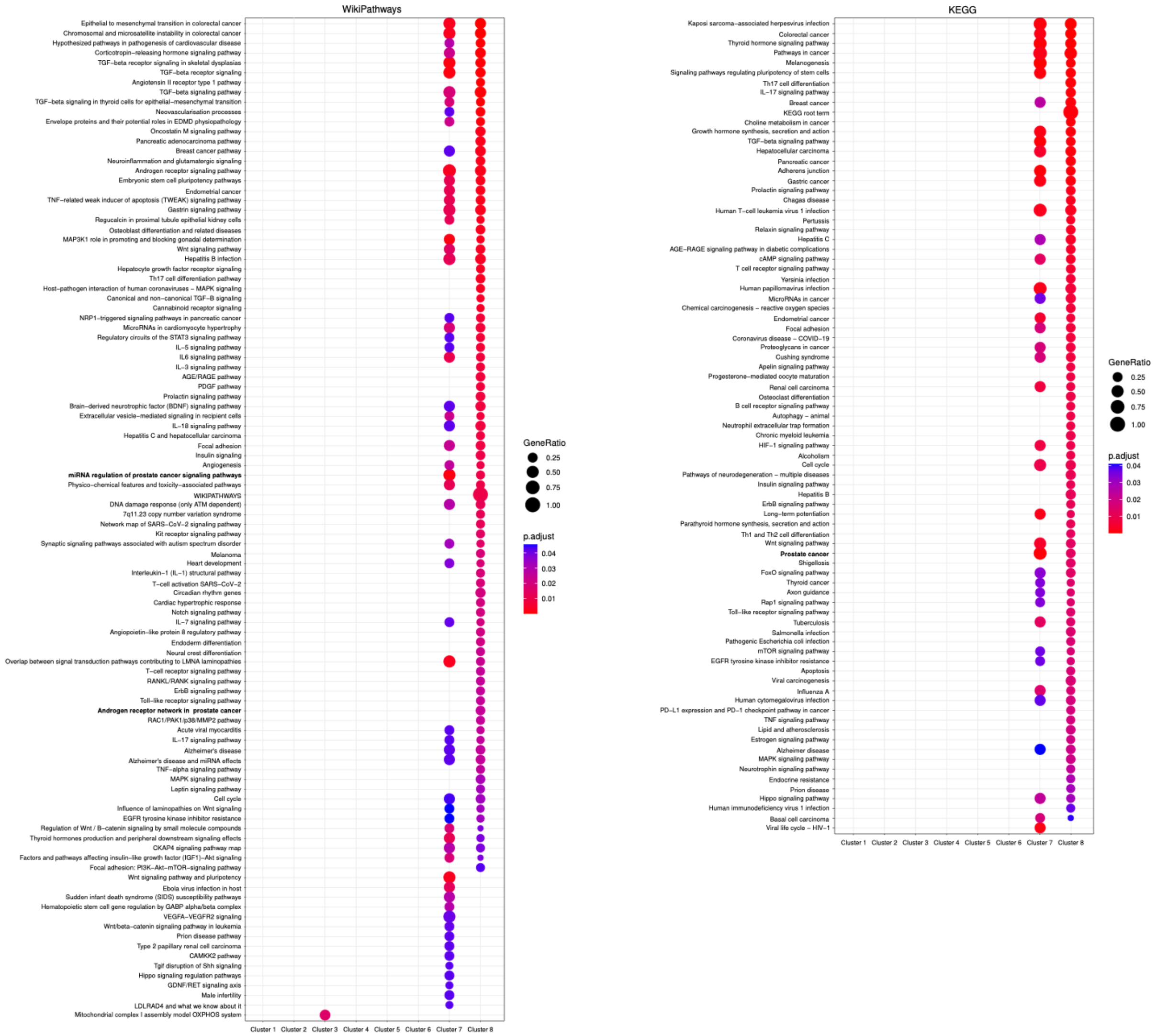


**Figure S8. Functional enrichment analysis using g:Profiler to compare the central proteins across different clusters.** Two databases of pathways were interrogated, WikiPathways (left [4]), and KEGG (right [5]). Overall clusters 1, 2, 3, 4, and 6 did not show any enrichments, possibly due to their higher number of central proteins compared to clusters 5, 7, and 8. Among the clusters with enrichments, only cluster 7 showed similarities to cluster 8, such as enrichment in *prostate cancer* (adjusted p-value = 2.0e-2). Cluster 8 also shows a significant prostate specific WikiPathway *Androgen receptor network in prostate cancer*.

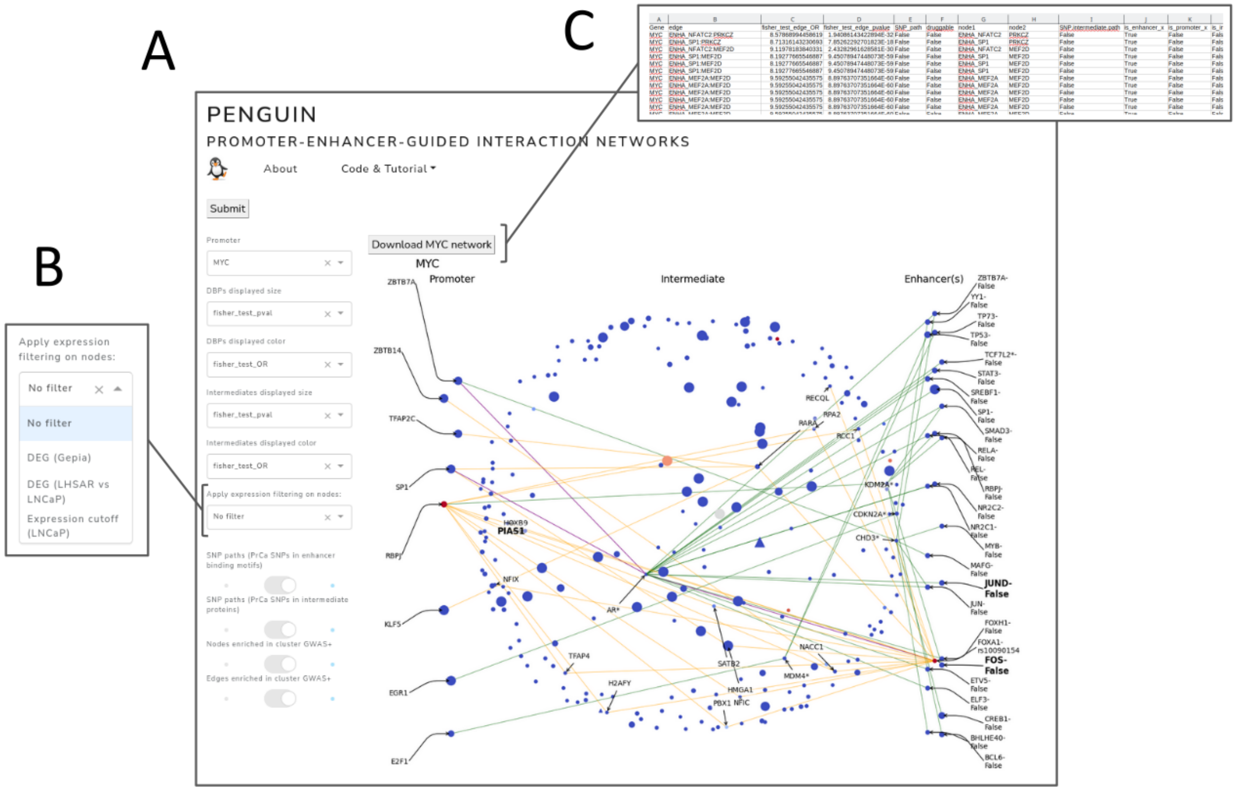


**Figure S9. PENGUIN web server. (A)** Screenshot of the main page of the PENGUIN web server where the EPIN of the *MYC* promoter is visualized with SNP paths highlighted (orange, PrCa SNPs in enhancer binding motifs; green, PrCa SNPs in intermediate proteins; purple, both). **(B)** Example of displaying node filtering options based on gene expression (deregulated genes, DEGs, identified using Gepia resource, LHSAR versus LNCaP). **(C)** Option to download network and associated statistics for each of the over 4 thousands PrCa EPINs available.

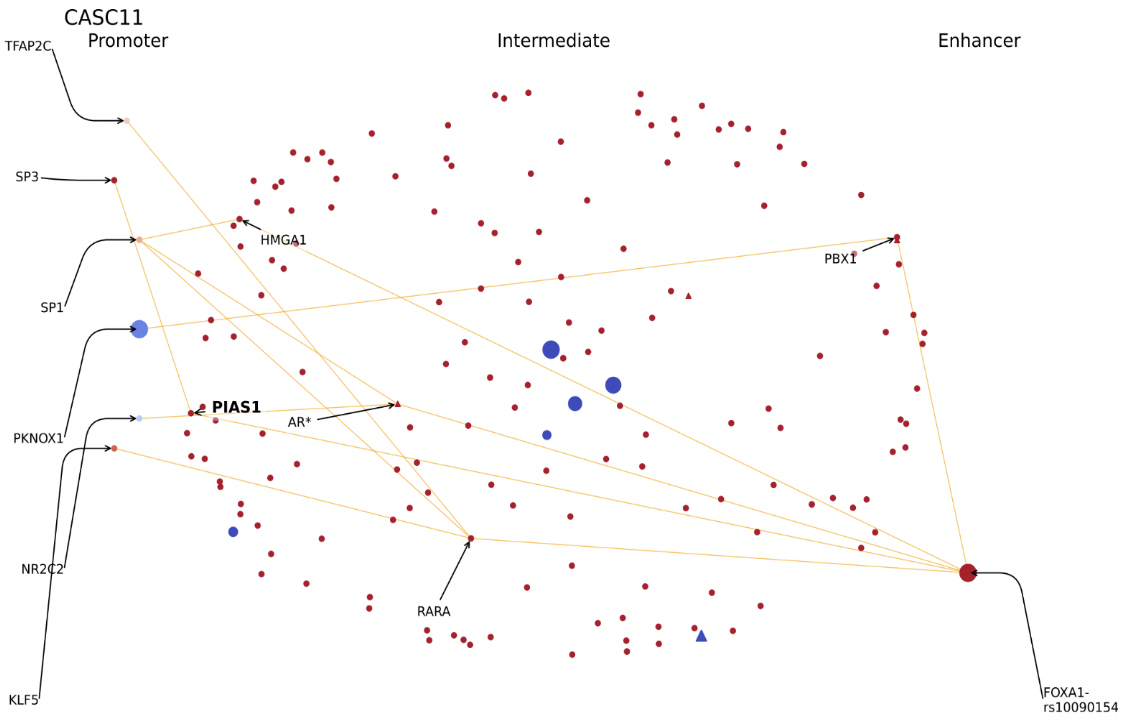


**Figure S10A. The CASC11 example.** The EPIN of CASC11 promoter is affected by variant rs10090154, the same well known variant associated with risk of developing prostate carcinoma that we introduced with MYC EPIN. The promoter binds 6 proteins: TFAP2C, SP3, SP1, PKNOX1, NR2C2 and KLF5. Potentially affected protein interactors of the EPIN include: HMGA1, PIAS1, AR, RARA, and PBX1. The yellow lines represent the set of edges that bridge promoter-bound DBPs and intermediate proteins with enhancer-bound DBPs with PrCa SNPs falling in their binding motif

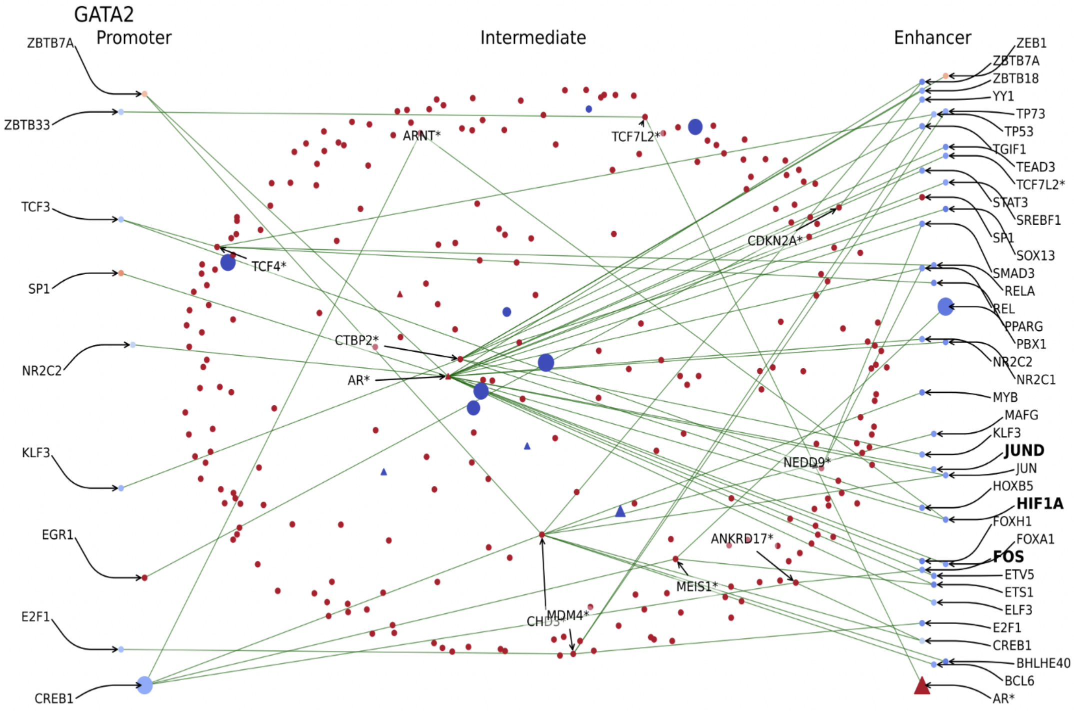


**Figure S10B. The GATA2 example.** GATA2 EPIN presents up to 11 intermediates affected by PrCa related SNPs, namely TCF4, CTBP2, AR, ARNT, TCF7L2, CDKN2A, NEDD9, ANKRD17, MEIS1, MDM4 and CHD3.Proteins bound to the promoter region include: ZBTB7A, ZBTB33, TCF3, SF1, NR2C2, KLF3, EGR1, E2F1 and CREB1, but most importantly, the EPIN presents AR bound to the enhancer region, which, as we pointed out with MYC, is the target of several PrCa drugs. The green lines represent the set of edges that bridge promoter-bound DBPs with enhancer-bound DBPs through intermediate proteins with PrCa SNPs falling in the genomic region of the corresponding coding gene.

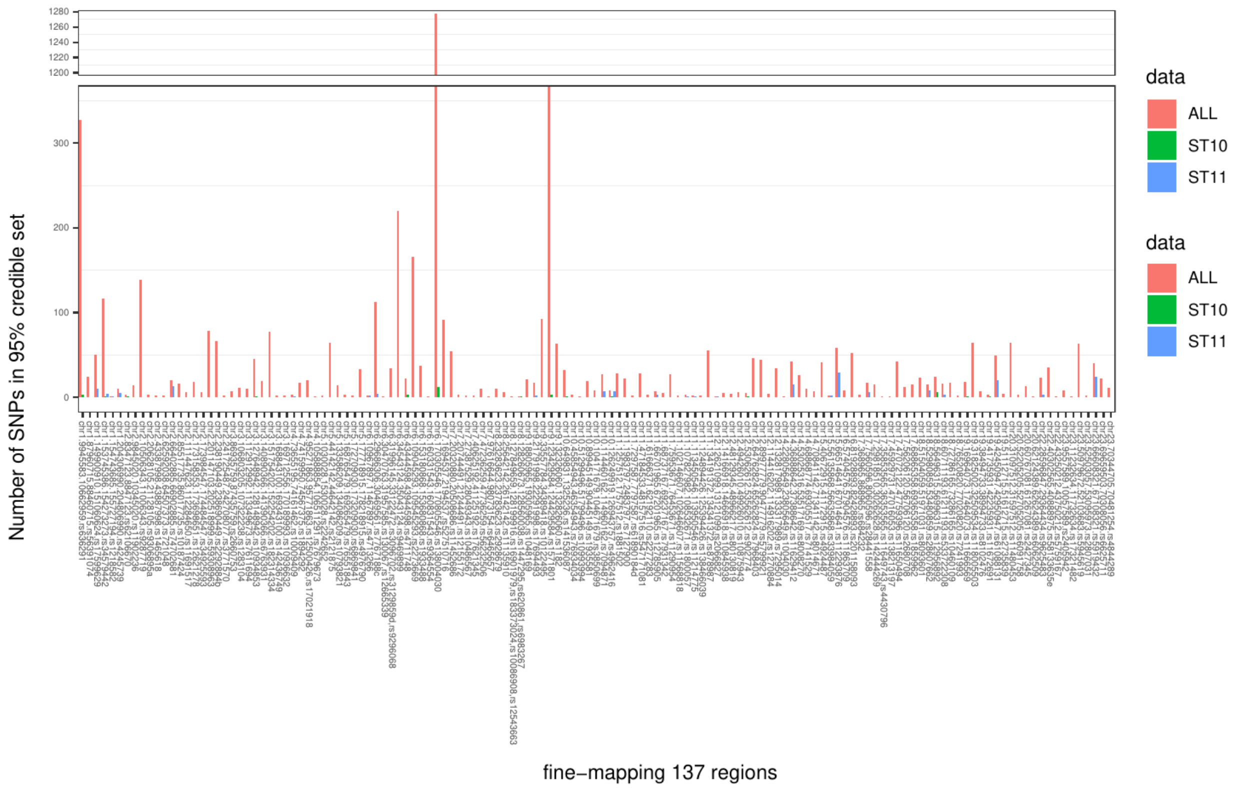


**Figure S11. Fine mapped regions.** x-axis illustrates 137 regions previously associated with PrCa; y-axis the number of PrCa SNPs (95% credible set) in each region, across ALL (5,412 PrCa SNPs) in red. Color-coded parallel bars in green and blue illustrate the location of the PrCa SNPs identified in ST10 and ST11 and characterized by PENGUIN. No significant correlation (Pearson r=0.2, p-value=0.06 and Pearson r=0.1, p-value=0.3, for ST10 and ST11, respectively) was identified between the number of PrCa SNPs in the regions and the number of PrCa SNPs we prioritized in this work.

### Tables

***Table S1. Information on 4,314 EPIN Promoters.***

Each row contains a unique promoter, which we call EPIN promoter. Each EPIN promoter: is exclusive to only one cluster; can have multiple enhancer elements linked to it by E-P loop; can have proteins linked to it by PPI in the network.

***EPIN_promoter****: Gene name (RefSeq hg19 from UCSC Genome Browser)*
***EPIN_promoter_anchor****: genomic positions of the promoter (hg19)*
*enhancers: genomic positions of the enhancer (hg19)*
***GWAS_overlap****: TRUE/FALSE indicating whether the EPIN enhancer(s) overlap PrCa SNP*
***CTCF_overlap****: TRUE/FALSE indicating whether the promoter regions AND EPIN enhancer(s) overlap CTCF binding site*
***Oncogene_overlap****: TRUE/FALSE indicating whether the EPIN promoter is an oncogene (Methods)*
***n_enha****: Number of HiCHip defined non-prioritized enhancers of the EPIN loop*
***n_prioritized_enha****: Number of prioritized enhancers of the EPIN loop*
***Cluster****: cluster number that The EPIN belongs*
***CTCF****: CTCF enrichment estimated by Fisher test, of the cluster that this EPIN belongs to. Can be either +/-/=.*
*GWAS: GWAS paintor enrichment estimated by Fisher test, of the cluster that this EPIN belongs to. Can be either +/-/=.*
***GWAS_Cat_prostate****.carcinoma; GWAS Catalogue (prostate carcinoma) enrichment estimated by Fisher test, of the cluster that this EPIN belongs to. Can be either +/-/=*
***max_PET_Q0.01****: HiChIP max q score*
***min_Q.Value_Bias****: the Q value is the pvalue from the loops in FitHiChIP*
***oncogene****: Whether the EPIN promoter gene is a previously reported oncogene*
***LNCaP****: FPKM cell-specific gene expression. Average of two replicates(LNCaP_1, LNCaP_2)*
***DE_gepia_EPIN_Promoter****: Differential gene expression of EPIN_Promoter from Gepia (see Methods Differential Gene Expression)*
***DE_LNCaP_LHSAR_EPIN_Promoter****: Differential gene expression of EPIN_Promoter from LNCaP versus LHSAR cell lines (see Methods Differential Gene Expression)*
***n_intermediate..no.DBP.*** *Number of Intermediate nodes in the PPI of the EPIN excluding the DBP bound to the E/P anchors*
***n_intermediate_with_SNP_geneLoc****: Number of n_intermediate..no.DBP with a SNP (GWAS paintor) overlapping their Genomic location.*
***prop_intermediate_with_SNP_geneLoc****: The proportion: n_intermediate_with_SNP_geneLoc / n_intermediate..no.DBP*
***n_DE_intermediate_gepia****: Number of n_intermediate..no.DBP that are DE from Gepia (see Methods Differential Gene Expression)*

***N_intermediate_in_deseq**: Number of n_intermediate..no.DBP that are present in deseq list ((see Methods Differential Gene Expression)*
***n_DE_intermediate_deseq**: Number of n_intermediate..no.DBP that are DE in the deseq list (see Methods Differential Gene Expression)*
***prop_DE_intermediate_deseq**: the proportion: n_DE_intermediate_deseq / N_intermediate_in_deseq*
***enrichment_DE_deseq_SNP.bs.DBP.path**: Enrichment of DE genes by deseq in the PPI paths defined from a DBP binding site that overlaps with a SNP (GWAS paintor). (see Methods Differential Gene Expression, last paragraph)*
***enrichment_DE_deseq_SNP.intermediate.path**: Enrichment of DE genes by deseq in the PPI paths defined by intermediates that have a SNP (GWAS paintor) overlapping their Gene Genomic location.*

***Table S2.** List of 885 unique proteins that are nodes among the 4314 EPIN_promoter. Three types of nodes exist: with DNA binding motifs in the promoter (promoter-bound nodes), proteins with DNA binding motifs in the enhancers (enhancer-bound nodes), and proteins interacting with promoter-bound or enhancer-bound nodes but without DNA binding motifs onto the promoter or the enhancers (intermediate nodes). 751 out of intermediate and as DNA-bound nodes in different EPINs. 261 unique DNA-binding proteins have predicted binding sites in at least one of the anchors of enhancers and promoters. We annotated as PrCa druggable each protein node that is target for drugs that are assigned as Approved or under Clinical Trials (Phase 1, 2, 3) or Investigable for Prostate Cancer.*

***Gene.name**: corresponding Gene name of the encoded protein*
***Is_intermediate_node**: Whether this Gene is an intermediate node in the PPI network of an EPIN*
***is_DBP_node**: Whether this Gene is a DBP protein in the E/P anchors*
***DrugBank_drugs**: DrugBank drugs for targeting that Gene*
***PrCa_drugs**: DrugBank drugs, specific for PrCa for targeting that Gene*
***DE_in_gepia**: If the Gene is DE from Gepia (see Methods)*
***DE_in_deseq**: If the Gene is DE from deseq (see Methods)*
***LNCaP_1**: cell-specific gene expression. replicates(LNCaP_1, LNCaP_2)*
***LNCaP_2**: cell-specific gene expression. replicates(LNCaP_1, LNCaP_2)*
***Oncogene**: Indicates whether that gene is reported as a known oncogene*

***Table S3A. General characteristics of the eight identified chromatin cluster.***

***cluster**: Cluster number ID*
***N_networks**: Number of EPIN_promoters belonging to the cluster.*
***Mean_enhancers**: The mean number of prioritized enhancers per EPIN_promoter belonging to that cluster.*
***Sd_enhancers**: The standard deviation of the number of prioritized enhancers per EPIN_promoter belonging to that cluster.*

***sd_betweenness****: The standard deviation of the mean betweennesses of the PPI network per EPIN_promoter belonging to that cluster.*
***mean_betweenness****: The average of the mean betweennesses of the PPI network per EPIN_promoter belonging to that cluster.*
***sd_degree****: The standard deviation of the mean degree of the PPI network per EPIN_promoter belonging to that cluster.*
***mean_degree****: The average of the mean degree of the PPI networks per EPIN_promoter belonging to that cluster.*
***mean_edges****: The average of the number of edges of the PPI networks per EPIN_promoter belonging to that cluster.*
***sd_edges****: The standard deviation of the number of edges of the PPI networks per EPIN_promoter belonging to that cluster.*
***mean_P_binders****: The average Promoter binders per EPIN_promoter belonging to that cluster.*
***sd_P_binders****: The standard deviation of Promoter binders per EPIN_promoter belonging to that cluster.*
***mean_E_binders****: The average Enhancer binders per EPIN_promoter belonging to that cluster.*
***sd_E_binders****: The standard deviation of the Enhancer binders per EPIN_promoter belonging to that cluster.*
***mean_intermediates****: The average intermediate nodes per EPIN_promoter belonging to that cluster.*
***sd_intermediates****: The standard deviation of the intermediate nodes per EPIN_promoter belonging to that cluster.*
***n_enriched_edges****: Number of enriched edges of that cluster estimated by Fisher test compared to all other clusters. (see Methods)*
***N_enriched_druggable_edges****: Number of enriched edges with druggable indication of that cluster estimated by Fisher test.(see Methods)*
***N_oncogene_networks****: Number of EPIN promoter genes previously reported as known oncogenes.*
***OR_oncogenes****: OddRatio estimated by Fisher test of the enrichment of previously reported known oncogenes in that cluster.*
***pv_oncogenes****: p-value estimated by Fisher test of the enrichment of previously reported known oncogenes in that cluster.*

***CTCF****: CTCF peak enrichment estimated by Fisher test. Can be either +/-/=.*
***CTCF_OR****: OddRatio estimated by Fisher test of the enrichment of CTCF peak binding in the EPINs belonging to the cluster*
***CTCF_pv****: p-value estimated by Fisher test of the enrichment of CTCF binding in the EPINs belonging to the cluster*

***GWAS****: GWAS paintor enrichment estimated by Fisher test. Can be either +/-/=.*
***GWAS_OR****: OddRatio estimated by Fisher test of the enrichment of GWAS paintor SNPs in the EPINs belonging to the cluster*
***GWAS_pv****: p-value estimated by Fisher test of the enrichment of GWAS paintor SNPs in the EPINs belonging to the cluster*

***GWAS_Cat_PrCa***; *GWAS Catalog (prostate carcinoma) SNPs enrichment estimated by Fisher test. Can be either +/-/=.*
***GWAS_Cat_PrCa_OR***: *OddRatio estimated by Fisher test of the enrichment of GWAS Catalog (prostate carcinoma) SNPs in the EPINs belonging to the cluster*
***GWAS_Cat_PrCa_pv***: *p-value estimated by Fisher test of the enrichment of GWAS Catalog (prostate carcinoma) SNPs in the EPINs belonging to the cluster*
***EPIN_promoter_LNCaP_mean_expression***: *Mean of cell-specific gene expression of the EPINs of that cluster*
***EPIN_promoter_LNCaP_sd_expression***: *Standard deviation of cell-specific gene expression of the EPINs of that cluster*
***EPIN_promoter_n_gepia_diff_exp_neworks***: *Number of EPIN promoter Genes that are DE from Gepia dataset.*
***EPIN_promoter_n_lncap_lhsar_diff_exp_neworks***: *Number of EPIN promoter Genes that are DE in LNCaP vs LHSAR*

***Table S3B:*** *Clustering Comparison for LNCaP and LHSAR cell lines*

***Cell_line***: *The Cell Line*
***Cluster_num***: *Total number of clusters to cut the hierarchical clustering Tree*
***Cluster***: *The cluster number ID*

***CTCF_ENCFF155SPQ***: *Enrichment of CTCF ChiPseq peaks from ENCODE for LNCaP Cell Line in the cluster. +/-/= as estimated with Fisher Test.*
***CTCF_ENCFF155SPQ_OR***: *Enrichment of CTCF ChiPseq peaks from ENCODE for LNCaP Cell Line peaks in the cluster. OddRatio as estimated with Fisher Test.*
***CTCF_ENCFF155SPQ_pv***: *Enrichment of CTCF ChiPseq peaks from ENCODE for LNCaP Cell Line peaks in the cluster. P-value as estimated with Fisher Test.*

***CTCF_SRX3322103.10***: *Enrichment of CTCF ChiPseq from SRX3322103 at q-value of 1e-10 for Prostate Epithelial Cells. +/-/= as estimated with Fisher Test.*
***CTCF_SRX3322103.10_OR***: *Enrichment of CTCF ChiPseq from SRX3322103 at q-value of 1e-10 for Prostate Epithelial Cells. OddRatio as estimated with Fisher Test.*
***CTCF_SRX3322103.10_pv***: *Enrichment of CTCF ChiPseq from SRX3322103 at q-value of 1e-10 for Prostate Epithelial Cells. P-value as estimated with Fisher Test.*

***CTCF_SRX3322104.10***: *Enrichment of CTCF ChiPseq from SRX3322104 at q-value of 1e-10 for Prostate Epithelial Cells. +/-/= as estimated with Fisher Test.*
***CTCF_SRX3322104.10_OR***: *Enrichment of CTCF ChiPseq from SRX3322104 at q-value of 1e-10 for Prostate Epithelial Cells. OddRatio as estimated with Fisher Test.*
***CTCF_SRX3322104.10_pv***: *Enrichment of CTCF ChiPseq from SRX3322104 at q-value of 1e-10 for Prostate Epithelial Cells. P-value as estimated with Fisher Test.*

***CTCF_SRX332210_3_4.10***: *Enrichment of CTCF ChiPseq from concatenation of SRX3322104 and SRX3322103 at q-value of 1e-10 for Prostate Epithelial Cells. +/-/= as estimated with Fisher Test.*
***CTCF_SRX332210_3_4.10_OR***: *Enrichment of CTCF ChiPseq from concatenation of SRX3322104 and SRX3322103 at q-value of 1e-10 for Prostate Epithelial Cells. OddRatio as estimated with Fisher Test.*
***CTCF_SRX332210_3_4.10_pv***: *Enrichment of CTCF ChiPseq from concatenation of SRX3322104 and SRX3322103 at q-value of 1e-10 for Prostate Epithelial Cells. P-value as estimated with Fisher Test.*

***GWAS***: *GWAS paintor enrichment estimated by Fisher test. Can be either +/-/=.*
***GWAS_OR***: *OddRatio estimated by Fisher test of the enrichment of GWAS paintor SNPs*
***GWAS_pv***: *p-value estimated by Fisher test of the enrichment of GWAS paintor SNPs*

***GWAS_Cat_prostate.carcinoma***: *GWAS Catalog (prostate carcinoma) SNP enrichment estimated by Fisher test. Can be either +/-/=*
***GWAS_Cat_prostate.carcinoma_OR***: *OddRatio estimated by Fisher test of the enrichment of GWAS Catalog (prostate carcinoma) SNP*
***GWAS_Cat_prostate.carcinoma_pv***: *p-value estimated by Fisher test of the enrichment of GWAS Catalog (prostate carcinoma) SNPs*

***CTCF_FIMO***: *Enrichment of CTCF binding sites predicted by FIMO. +/-/= as estimated with Fisher Test.*
***CTCF_FIMO_OR***: *Enrichment of CTCF binding sites predicted by FIMO. OddRatio as estimated with Fisher Test.*
***CTCF_FIMO_pv***: *Enrichment of CTCF binding sites predicted by FIMO. P-value as estimated with Fisher Test.*

***total_genes:*** *Number of genes in the cluster*
***total_oncogenes:*** *Number of genes annotated as previously reported oncogenes in the cluster*
***oncogenes_pval:*** *p-value estimated by Fisher test of the enrichment of previously reported known oncogenes in that cluster.*
***oncogenes_OR:*** *OddRatio estimated by Fisher test of the enrichment of previously reported known oncogenes in that cluster.*

***GWAS_non_fine_mapped:*** *Enrichment of non-fine mapped list of PrCa GWAS SNPs. +/-/= as estimated with Fisher Test.*
***GWAS_non_fine_mapped_OR:*** *OddRatio estimated by Fisher test of the enrichment of non-fine mapped list of PrCa GWAS SNPs.*
***GWAS_non_fine_mapped_pv:*** *p-value estimated by Fisher test of the enrichment of non-fine mapped list of PrCa GWAS SNPs.*

***Table S4: Enrichment of PrCa-relevant annotations in cluster 8 when including and excluding intermediates.***

*EPIN reconstruction for 0 (DBP - DBP direct PPI interaction) and 1 intermediate (DBP - intermediate - DBP PPI interaction)*
*Enrichment of Oncogenes, CTCF overlap and GWAS paintor SNP overlap in the cluster that is GWAS and CTCF enriched.*
***OR**: OddRatio estimated by Fisher*
***pvalue**: p-value estimated by Fisher*

***Table S5: Cluster 8 pathways.***
*Adapted from gprofiler2 R package documentation:*
*https://cran.r-project.org/web/packages/gprofiler2/vignettes/gprofiler2.html#enrichment-analysis*
***query**: the name of the input query variable with the input genes.*
***significant**: Boolean indicating statistical significance.*
***p_value**: g:Profiler GSEA hypergeometric test p-value (Benjamini-Hochberg FDR Correction)*
***term_size**: number of genes that are annotated to the term*
***query_size**: number of genes that were included in the query. This might be different from the size of the original list if any genes failed to be mapped to Ensembl gene IDs or if the gene has no annotation in the corresponding database.*
***intersection_size**: the number of genes in the input query that are annotated to the corresponding term*
***precision**: the proportion of genes in the input list that are annotated to the function (defined as intersection_size/query_size)*
***recall**: the proportion of functionally annotated genes that the query recovers (defined as intersection_size/term_size)*
***term_id**: unique term identifier (e.g GO:0005005) from the corresponding target database*
***source** : ID of the target database for g:Profiler GSEA.*

*IDs:*
*GO:BP -> Gene Ontology Biological Process*
*GO:MF -> Gene Ontology Molecular Function*
*GO:CC -> Gene Ontology Cellular Component*
*KEGG -> KEGG Pathways*
*REAC -> Reactome Pathways*
*WP -> WikiPathways*
*TF -> TRANSFAC*
*MIRNA -> miRTarBase*
*HPA -> Human Protein Atlas*
*CORUM -> CORUM comprehensive resource of mammalian protein complexes*
*HP -> Human Phenotype Ontology (HPO)*

***term_name**: Tested database entry name*
***effective_domain_size**: the total number of genes “in the universe” used for the hypergeometric test*
***source_order**: numeric order for the term within its data source (this is important for drawing the results)*
***parents**: list of term IDs that are hierarchically directly above the term. For non-hierarchical data sources this points to an artificial root node.*
***evidence_codes**: a lists of all evidence codes for the intersecting genes between input and the term. The evidences are separated by comma for each gene.*

***intersection****: a comma separated list of genes from the query that are annotated to the corresponding term*

***Table S6:*** *Enrichment from testing GWAS signals across 17 diseases in the eight clusters identified by PENGUIN (SNP assigned to cluster/trait compared to other cluster/trait). GWAS SNPs are obtained from GWAS Catalog (Method).*

***TRAIT****: Name of trait from GWAS data analyzed*
***SNPS****: Number of SNPs from GWAS data analyzed*
***Cluster****: Cluster from PENGUIN*
***N_in_cluster****: Number of EPINs overlapping the SNPs form that trait inside the cluster (used for Fisher tests)*
***N_out_cluster****: : Number of EPINs overlapping the SNPs form that trait outside of the cluster (used for Fisher tests)*
***Total_n_in_cluster****: Total number of EPINs in the cluster*
***Total_n_out_cluster****: Total number of EPINs outside the cluster*
***N_in_cluster_no_disease****: Number of EPINs inside the cluster that dont overlap the SNPs form that trait (used for Fisher tests)*
***N_out_cluster_no_diseasep****: Number of EPINs outside the cluster that dont overlap the SNPs form that trait (used for Fisher tests)*
***P****: p-value of enrichment from Fisher test (Methods)*
***OR****: odds ratio of enrichment from Fisher test (Methods)*
*gap: distance*

***Table S7:*** *Intermediate protein pathways.*
*Same as Table S5 and Table S9.*

***Table S8:*** *Intermediate Protein Centrality significance.*
***all_genes****: Protein gene ID (HGNC)*
***mean_bet_in****: Mean of node betweenness centrality of the intermediate nodes in EPINs belonging to Cluster 8.*
***mean_bet_out****: Mean of node betweenness centrality of the intermediate nodes in EPINs belonging to Clusters 1 to 7.*
***ratio_bet****: Ratio between the mean node betweenness centrally inside and outside of Cluster 8. Equals to mean_bet_in / mean_bet_out.*
***mean_deg_in****: Mean of node degree centrality of the intermediate nodes in EPINs belonging to Cluster 8.*
***mean_deg_out****: Mean of node degree centrality of the intermediate nodes in EPINs belonging to Clusters 1 to 7.*
***ratio_deg****: Ratio between the mean node degree betweenness centrally inside and outside of Cluster 8. Equals to mean_deg_in / mean_deg_out .*
***pvalue_ratio****: Statistical significance of the node betweenness ratio (ratio_bet) (Methods: Enriched intermediate nodes within each cluster). p-value represents the probability of finding the*

*protein betweenness specificity ratio to be higher or equal to the real betweenness ratio value upon random network cluster generation.*
***p-value_degree**: Statistical significance of the node degree ratio (ratio_deg) (Methods: Enriched intermediate nodes within each cluster). p-value represents the probability of finding the protein degree specificity ratio to be higher or equal to the real degree specificity ratio value upon random network cluster generation.*

***Table S9:** 22 intermediate proteins*
*Same as Table S5 and Table S7.*

***Table S10:** SNP paths TF. 36 PrCa SNPs falling within 60 DBP motifs in enhancer regions linking 34 different promoters whose EPINs include 5,184 edges. Among these, we identified 17 PrCa SNPs falling within 16 promoter EPINs (1,894 edges) belonging to the GWAS+ cluster that had at least one PrCa SNP in their enhancers.*

***SNP**: unique rsID of PrCa SNP falling within enhancer DNA binding motifs.*
***EPIN_promoter**: Gene name (same as Table S1)*
***Cluster**: same as Table S1*
***DBP**: The DBP Gene name that overlaps with the SNP*
***intermediates (also DBP)**: Gene names of the PPI subnetwork starting from the DBP bound to the enhancer anchor with the SNP overlapping the DNA binding motif and including all intermediate nodes interacting with it.*
***n_intermediates (also DBP)**: Number of nodes (intermediates (also DBP)*
***PVAL_PrCa**: p-value associated with the SNP overlapping the binding site of the DNB.*

***diffexpr_DBP.gepia.gepia**: DE of the DBP from Gepia dataset (see Methods Differential Gene Expression)*
***diffexpr_DBP.lncap_lhsar.gepia**: DE of the DBP from Gepia dataset (see Methods) comparing LNCaP and LHSAR cell lines (see Methods Differential Gene Expression)*

***EPIN_promoter_log2FC.gepia**: DE (log2FC) of the DBP from Gepia dataset (see Methods Differential Gene Expression)*
***EPIN_promoter_adjp.gepia**: DE (adjusted p-value) of the DBP from Gepia dataset (see Methods Differential Gene Expression)*
***Numb_diffexpr_unique_intermediate_pathways.gepia**: Number of nodes in the PPI subnetwork (including intermediate nodes and DBP that are DE from Gepia dataset.*
***Perc_diffexpr.gepia**: the percentage of Numb_diffexpr_unique_intermediate_pathways.gepia / n_intermediates (also DBP)*
***Diffexpr_intermediate_pathways.gepia**: The nodes in the PPI subnetwork (including intermediate nodes and DBP that are DE from Gepia dataset (see Methods Differential Gene Expression)*

***EPIN_promoter_log2FC.lncap_lhsar**: The DE log2FC of LNCaP vs LHSAR for the EPIN promoter Gene (see Methods Differential Gene Expression)*

***EPIN_promoter_adjp.lncap_lhsar**: The p-value adjusted for the DE of LNCaP vs LHSAR for the EPIN promoter Gene dataset (see Methods Differential Gene Expression)*
***Numb_diffexpr_unique_intermediate_pathways.lncap_lhsar**: Number of nodes in the PPI subnetwork (including intermediate nodes and DBP) that are DE in LNCaP vs LHSAR (see Methods Differential Gene Expression)*

***Perc_diffexpr.lncap_lhsar**: The percentage of Numb_diffexpr_unique_intermediate_pathways.lncap_lhsar / n_intermediates (also DBP)*

***Diffexpr_intermediate_pathways.lncap_lhsar**: The nodes in the PPI subnetwork (including intermediate nodes and DBP that are DE in LNCaP vs LHSAR (see Methods Differential Gene Expression)*

***EPIN_promoter_pval_pqtl**: lists the pQTL p-value of the association of the SNP with the protein levels encoded by the EPIN_promoter*
***EPIN_promoter_seqids_pqtl**: lists the identity of the proteins by SeqIDs for the association of the SNP with the protein levels encoded by the EPIN_promoter*
***Intermediate_pathways_pval_pqtl**: lists the pQTL p-value of the association of the SNP with the protein levels encoded by the proteins in intermediates*
***Intermediate_pathways_seqids_pqtl**: lists the pQTL p-value of the association of the SNP with the protein levels encoded by the proteins in intermediates*
***pqtl_proteins**: lists the identity of the proteins by SeqIDs for the association of the SNP with the protein levels encoded by the proteins in intermediates*

***Table S11:** Intermediates with SNPs. Network paths link 172 PrCa SNPs falling within 26 gene bodies coding for intermediate proteins, of which 7 are known oncogenes (MAP2K1, CHD3, AR, SETDB1, ATM, CDKN1B, USP28). We identify edges that are most enriched in our GWAS+ cluster which could be pointing to essential links between the gene encoding for the intermediate and containing a PrCa predisposing SNP at a particular EPIN.*
***SNP**: SNP rsID*
***gene_w_SNP**: The gene name of the intermediate node that has the SNP in it's genomic location*
***PVAL_PrCa**: P-value associated with the SNP*
***gene_w_SNP_log2FC.gepia**: The DE log2FC from Gepia dataset of the Gene that has the SNP in it's genomic location (see Methods Differential Gene Expression)*
***gene_w_SNP_adjp.gepia**: The DE p-value adjusted from Gepia dataset of the Gene that has the SNP in it's genomic location (see Methods Differential Gene Expression)*
***EPIN_promoters_counts**: Number of EPIN promoters that the intermediate node with the SNP in its genomic location is present*
***Clusters**: The clusters that contain the EPIN_promoters_counts*

***EPIN_promoters_top_DE_Gepia:*** *Top 3 EPIN promoters from EPIN_promoters_counts, ranked by DE from Gepia. Columns show the corresponding DE LogFC values. (see Methods Differential Gene Expression)*
***EPIN_promoters_top_DE_pval_Gepia:*** *Top 3 EPIN promoters from EPIN_promoters_counts, ranked by DE from Gepia. Columns show the corresponding DE p-values. (see Methods Differential Gene Expression)*
***EPIN_promoters_DE_Gepia:*** *Number of EPIN promoters from EPIN_promoters_counts that are DE from Gepia. (see Methods Differential Gene Expression)*

***EPIN_promoters_top_DE_LNCAP_LHSAR:*** *Top 3 EPIN promoters from EPIN_promoters_counts, ranked by DE from LNCaP vs LHSAR analysis. Columns show the corresponding LogFC values. (see Methods Differential Gene Expression)*
***EPIN_promoters_top_DE_pval_LNCAP_LHSAR:*** *Top 3 EPIN promoters from EPIN_promoters_counts, ranked by DE from LNCaP vs LHSAR analysis. Columns show the corresponding p-values. (see Methods Differential Gene Expression)*
***EPIN_promoters_DE_LNCAP_LHSARcount:***

***EPIN_promoters_pval_pqtl_genes:*** *[ lists the proteins encoding for a gene among the EPIN_promoters with a p-value of the association < 10^-4 ] -> i can't remmeber if this was association with any SNP or with the particular SNP listed*
***gene_w_SNP_pval_pqtl_genes:*** *lists the pQTL p-value of the association of the SNP with the protein levels encoded by the gene_w_SNP*
***EPIN_promoters_pval_pqtl_genesminp:*** *lists among all the genes among the EPIN_promoters the gene/protein with the lowest p-value of association*
***EPIN_promoters_pval_pqtlminp:*** *lists the p-value of association with the EPIN_promoter*

***gene_w_SNP_DrugBank_drugs:*** *DrugBank drugs for targeting the gene_w_SNP*
***gene_w_SNP_PrCa_drugs:*** *DrugBank drugs, specific for PrCa for targeting the gene_w_SNP*
***gene_w_SNP_is_oncogene:*** *Whether the gene_w_SNP is reported as previously known gene*
***sig***
***edge:*** *Most enriched edge (estimated by Fisher test) passing through gene_w_SNP*
***fisher_test_edge_OR:*** *Column reporting the maximum enrichment's OddsRatio, estimated by Fisher test, among all clusters, of the most enriched edge passing through gene_w_SNP.*
***enriched_edge_cl8:*** *Column reporting the enrichment's OddsRatio for cluster 8, estimated by Fisher test, of the most enriched edge passing through gene_w_SNP.*

***Table S12:*** *number of PrCa SNPs in intermediates (Alex). We found that the GWAS+ cluster has the highest proportion of PrCa SNPs in the intermediate nodes compared to all other clusters (mean = 53.2, SE = 18.0, p-value <= 0.01, Table S12).*
***Cluster:*** *Cluster number ID*
***rsids per EPIN (mean)****: Number of PrCa SNPs in the intermediate nodes (mean among all EPINs of the cluster)*

***rsids per EPIN (sd):*** *Number of PrCa SNPs in the intermediate nodes (standard deviation among all EPINs of the cluster)*
